# Vegetative Phase Change Causes Age-Dependent Changes in Phenotypic Plasticity

**DOI:** 10.1101/2021.11.02.467012

**Authors:** Erica H. Lawrence-Paul, R. Scott Poethig, Jesse R. Lasky

**Affiliations:** Pennsylvania State University, Department of Biology, University Park, PA; University of Pennsylvania, Department of Biology, Philadelphia, PA

**Keywords:** age-dependent plasticity, developmental timing, ontogeny, phenotypic plasticity, plant development, vegetative phase change

## Abstract

- Phenotypic plasticity allows organisms to optimize traits for their environment. As organisms age, they experience diverse environments that benefit from varying degrees of phenotypic plasticity. Developmental transitions can control these age-dependent changes in plasticity and as such, the timing of these transitions can determine when plasticity changes in an organism.
- Here we investigate how the transition from juvenile-to adult-vegetative development known as vegetative phase change (VPC) contributes to age-dependent changes in phenotypic plasticity and how the timing of this transition responds to environment using both natural accessions and mutant lines in the model plant *Arabidopsis thaliana*.
- We found that the adult phase of vegetative development has greater plasticity in leaf morphology than the juvenile phase and confirmed that this difference in plasticity is caused by VPC using mutant lines. Further, we found that the timing of VPC, and therefore the time when increased plasticity is acquired, varies significantly across genotypes and environments.
- The consistent age-dependent changes in plasticity caused by VPC suggest that VPC may be adaptive. This genetic and environmental variation in the timing of VPC indicates the potential for population-level adaptive evolution of VPC.

## Introduction

Phenotypic plasticity, and its inverse – robustness -- are attributes of development in all organisms. When plasticity or robustness in a given trait is most beneficial for fitness depends on the degree of environmental variability, reliability in environmental cues and the costs associated with phenotypic adjustment (Fischer *et al*., 2014). Although there are many examples of adaptive plasticity (Ghalambor *et al*., 2007), we still have much to learn about what regulates plasticity, including whether plasticity changes in an organism’s lifecycle, which environmental cues bring about plasticity, and the drivers of genetic variation in plasticity.

As organisms age, they transition through different developmental phases that result in changing tradeoffs, while simultaneously experiencing diverse environments during those phases. As such, they might benefit from varying degrees of phenotypic plasticity across their lifespan. For example, cichlid fish display increased plasticity of morphological defenses (i.e. body size and coloration) in response to alarm cues just after birth and at the onset of reproduction when they are most vulnerable (Meuthen *et al*., 2018). Changes in plasticity associated with whole plant development are often overlooked (distinct from changes in plasticity associated with the development of a particular organ, e.g., changes in photosynthesis with leaf age (Niinemets, 2016)). A few studies demonstrate variation in plasticity before and after reproduction and at different phases of vegetative growth. For example, two different studies found that *Arabidopsis thaliana* and the aquatic plant *Sagittaria latifolia* have greater phenotypic plasticity in response to changes in nutrients post-flowering compared to pre-flowering (Zhang & Lechowicz, 1994; Dorken & Barrett, 2004). Older but pre-flowering *Plantago lanceolata* individuals show greater plasticity of chemical response to herbivory compared to younger individuals (Barton, 2008). In some cases, ontogenetic change in plasticity causes individuals of *different species* at the same developmental stage to respond more alike to environment than *conspecifics* at different developmental phases (Parrish & Bazzaz, 1985). Despite the significance of developmental changes in plasticity and their impacts on fitness in animals (Hoverman & Relyea, 2007; Fischer *et al*., 2014; Nilsson-Örtman *et al*., 2015; Meuthen *et al*., 2018; Sebestyén *et al*., 2020) we know relatively little about developmental changes in plasticity in plants.

In contrast, many studies examine plasticity in the timing of plant developmental transitions, mostly transitions from seed to seedling and vegetative to flowering. Plasticity in the timing of developmental transitions could allow plants to optimize their life history for their environment. For example, longer days and warmer temperatures accelerate flowering in *A. thaliana* (Levy & Dean, 1998; Blázquez *et al*., 2003) to promote flowering in the spring and summer when conditions are favorable, and seeds that mature during short photoperiodic days (i.e. autumn) are more sensitive to cold, delaying germination until after winter (Munir *et al*., 2001). Furthermore, the timing of developmental transitions and the degree of plasticity in other traits during developmental phases can be interconnected. For example, in *A. thaliana* a delay in flowering time was associated with greater plasticity in post-flowering traits (Zhang & Lechowicz, 1994). Together this indicates that the timing of developmental transitions can alter an organism’s phenotypic plasticity both by dictating which developmental phase the organism is in, and the degree of trait plasticity within that phase.

All plants transition between distinct juvenile and adult phases during vegetative development, which involves a wide range of changes in physiology and morphology. VPC is regulated by a highly conserved microRNA, miR156, and its targets, the Squamosa Promoter Binding-Like (SPL) transcription factors (Wu & Poethig, 2006; Willmann & Poethig, 2007; Wu *et al*., 2009; He *et al*., 2018). As individuals transition from a seedling to adult, changes in expression of the miR156/SPL module lead to phase specific differences in leaf morphology, photosynthetic traits, and growth strategies (Poethig, 1990; Bassiri *et al*., 1992; Bongard-Pierce *et al*., 1996; Telfer *et al*., 1997; Wang *et al*., 2011; Feng *et al*., 2016; Leichty & Poethig, 2019; Silva *et al*., 2019; Lawrence *et al*., 2020, 2021, 2022). Specifically, as plants transition from juvenile to adult, leaves become larger with decreased specific leaf area (SLA), and often display increased photosynthetic rates per unit leaf area. These changes in leaf morphology and physiology lead to a switch from a fast- to slow-growth strategy as plants transition from producing low-cost juvenile leaves to expensive adult leaves with long lifespans.

Because VPC alters how plants function, the types of structures produced, and the plant’s ability to transition to reproduction in some cases (Zhao *et al*., 2023), the timing of this transition is likely to have significant consequences for fitness. Further, the model presented in Fischer *et al*., (2014) suggests that age-dependent changes in plasticity are beneficial for fitness. Based on this model, an increase in plasticity with age would provide time early in life to collect enough information about the environment before investing in any costly phenotypic adjustment later in life. In contrast to reproduction and germination, very little is known about plasticity in the timing of vegetative phase change (VPC) or how it may impact the degree of phenotypic plasticity in other traits during vegetative development.

The timing of VPC is responsive to environmental cues, specifically light, and defoliation (Forster & Bonser, 2009; Yang *et al*., 2011; Leichty & Poethig, 2019; Rose *et al*., 2019; Xu *et al*., 2021). While the exact effects on the timing of VPC are unknown, expression of the miR156/SPL module is also altered by temperature, and drought (Kong *et al*., 2010; Lee *et al*., 2010; May *et al*., 2013; Arshad *et al*., 2018). Furthermore, the observation that juvenile and adult phases of alfalfa exhibit different tolerance to drought (Arshad *et al*., 2017) suggests any genotypic or species differences in the timing of this transition could be adaptive. However, the validity of this hypothesis is difficult to assess because the range of variation in the timing of VPC across species or even multiple genotypes within a species has only been investigated in a few cases.

Intimate relationships between development and plasticity led us to examine variation in the timing of VPC and the plasticity of phase-specific vegetative traits. In this study we examined eight transgenic lines with varying expression levels of the miR156/SPL module, to investigate how VPC impacts leaf morphological trait plasticity (i.e. age/phase-dependent plasticity) specifically, morphological traits that impact thermoregulation, hydraulic efficiency and light capture such as leaf serrations and overall leaf shape. We used these lines to further investigate how alterations in the timing of VPC impact plant growth responses across varying environments (i.e. heat, drought, low light), independent of genetic variation at other loci. Additionally, we used ten natural accessions of *A. thaliana* to investigate how much genetically regulated variation in the plasticity of developmental timing exists in this species, and whether relationships between VPC and plasticity observed in mutants is present across natural genotypes.

## Materials and Methods

### Plant growth and materials

Ten natural accessions of *Arabidopsis thaliana* (Table **S1**)—selected on the basis of a preliminary screen for variation in the timing of VPC—were obtained from the *Arabidopsis* Biological Resource Center (Ohio State University, Columbus, OH, USA). Accessions that transition earlier and later than the commonly used Col-0 background were selected, with the timing of VPC ranging from leaf 2 or 5.6 days to leaf 7.3 or 12.3 days, on average. We paired natural accessions with mutant lines in a Col-0 genetic background. Four of the mutant lines (*mir156a-2*, *mir156c-1*, *mir157c-1,* and *35S::MIM156)* accelerate the transition to the adult phase by reducing the levels of miR156 and miR157. The other four mutant lines (*35S::MIR156a*, *spl9-4, spl13-1* and the *MIR156A* genomic line) delay this transition. The *mir156a-2*, *mir156c-1*, *mir157c-1*, *35S::MIM156*, *35S::MIR156a*, and *spl9-4 spl13-1* lines have been previously described (Wu & Poethig, 2006; Franco-Zorrilla *et al*., 2007; Yang *et al*., 2013; Xu *et al*., 2016; He *et al*., 2018). The *MIR156A* genomic line developed in the Poethig lab is homozygous for a T-DNA insertion containing a 5.5 kb fragment spanning the region between the genes upstream and downstream of *MIR156A* in a pCAMBIA3301 backbone. These mutant and transgenic lines allowed us to evaluate developmentally juvenile and adult leaves at every leaf position within a single genetic background, thus distinguishing the effects of VPC from other factors such as length of exposure to the environment, and size or age of the plant, which could influence phenotypic plasticity.

Seeds were planted in 96 well flats in Fafard-2 growing mix supplemented with Peters 20-10-20 fertilizer. Beneficial nematodes (*Steinernema feltiae*, BioLogic, Willow Hill, PA), Marathon 1% granular insecticide and diatomaceous earth were added to the growing mix to control insects. Plants were placed in 4°C for 5 days before being grown in their respective treatment conditions (Table **S2**) in Percival growth chambers. Short day (10 hrs. light/14 hrs. dark, at 120 or 90 µmol m^-2^ s^-1^) light conditions were used to prevent flowering so adult vegetative phenotypes were not impacted by the reproductive transition. Although this prevents us from observing certain components of fitness such as fecundity, it ensures that observed developmental differences in plasticity are due to vegetative phase change. Plants were grown under 13.5W full spectrum LED lights. A reduction in light intensity for the low light treatment was achieved by increasing the distance between the lights and plants. Plants in all treatments were watered every other day with either 700 mL per flat until seeds germinated and throughout the 28-day growth period for control, heat and low light treatments or 350 mL per flat for drought treatment. Soil moisture was recorded with a Delta T HH2 Moisture Meter (Meter Group, Pullman, WA, USA) at a depth of approximately 1.5 cm twice a week both before and after watering at a minimum of two locations within the growth chamber. Temperature, light level, and average soil moisture conditions for each treatment are reported in Table **S2**

### Plant Phenotyping

Plants were harvested at the soil surface 28 days following transfer to growth chambers. Leaves were removed from the shoot and placed flat between two transparency sheets in the order they were initiated (leaf one is the first leaf to be produced) and scanned using a flatbed scanner (CanoScan LiDE220) at 300 dpi. All shoot material was then transferred to coin envelopes and dried at 60°C until constant mass.

Whole plant growth phenotypes were measured to determine the influence of the timing of VPC on plant productivity in the tested environments. Whole shoot mass from dried plants was recorded and shoot area (i.e. leaves and petioles) was analyzed from scanned images using FIJI (Fiji software). The number of leaves initiated was counted as the total number of fully expanded leaves and leaf primordia present at the end of the 28-day growing period. This value was used to calculate the average leaf initiation rate across the growing period by dividing the total number of leaves initiated by 28 days. Reductions in shoot mass, shoot area and number of leaves initiated as a result of being grown under environmental stress (i.e. drought, heat, and low light) were calculated as the difference between each genotype’s average value in control conditions and each individual grown in the stressed environment.

Individual leaf morphological traits were analyzed from scanned images using FIJI. Specifically, for leaf morphological measures, base leaf angle was measured as the angle between the edges of the leaf on either side of the petiole. Petioles were then removed, and the wand tool was used to select each leaf individually from binary images to measure area, perimeter, length, width and circularity and the presence or absence of serrations was noted. These traits describe both leaf shape and size to indicate phenotypic change across development and environment while also being reliably and easily measured from leaf scans.

Overall leaf shape was analyzed using morphometrics approaches in the Momocs package v1.4.0 for R (Bonhomme *et al*., 2014, R Core Team, 2021). Briefly, coordinates of outlines for individual leaf images were determined and elliptical fourier analysis was performed on the shapes using 20 harmonics and coefficients were normalized. Principal component analysis (PCA) was performed on the harmonic coefficients. Outlines from scanned leaves and the morphospace of the PCA (i.e. how leaf shape relates to PC values) can be viewed in the supplement (Fig. **S1**).

### Determination of Developmental Phase

Vegetative phase change in *Arabidopsis thaliana* is marked by changes in leaf morphology that include leaf base angle, length/width ratio, circularity, the presence and absence of serrations, and the presence or absence of abaxial trichomes (Telfer *et al*., 1997; Tsukaya *et al*., 2000; He *et al*., 2018). While the presence of abaxial trichomes is the most commonly used marker for the onset of the adult phase (Yang *et al*., 2013; Xu *et al*., 2016; He *et al*., 2018), we chose to use a combination of leaf morphological traits more easily measured from leaf scans to determine juvenile and adult phases. This method allows us to avoid the influence of any phase change-independent effects on trichome development (i.e. photoperiodic and UV-B effects on trichome regulators gibberelic acid biosynthesis and GLABRA3, respectively) that could arise in plants grown under different environments (Chien & Sussex, 1996; Olszewski *et al*., 2002; Yan *et al*., 2012). Further, by using a combination of traits we can get a more holistic representation of when vegetative phase change occurs, as the onset of different adult traits are often not perfectly synchronized.

Leaf stage was determined from leaf scans using a PCA summarizing phase specific traits (leaf base angle, the length:width ratio of the lamina, the presence vs. absence of serrations, and the circularity of the lamina), and comparing the PC1 value obtained by this approach to the PC1 value for plants over-expressing miR156 (35S::MIR156a, “OX”), in which all leaves are juvenile (Fig. **S2**). Leaves with a PC1 value greater than or equal to the value for the miR156-over expressing line were considered juvenile leaves, and leaves with a PC1 value less than this number were considered adult leaves. Specifically, the minimum PC1 value of all OX leaf samples for each treatment was identified. All leaves with PC1 values below this threshold were considered adult leaves. PC1 threshold values used here were 0.06 for control, 0.37 for drought, −1.91 for heat and 0.01 for low light conditions. Setting independent thresholds for each environment using leaves produced across all leaf positions eliminates the influence of any non-phase-change specific changes in PC1 values (i.e. plant size, age, or environment) in the determination of juvenile and adult leaves.

### Data Analysis and Statistics

Statistical analyses were performed in JMP ^®^ Pro v. 15.0.0 (SAS Institute Inc., Cary, NC). Differences in the time of transition among genotypes were compared by two-way ANOVA in where genotype and environmental treatment were the main effects. Additionally, GLMM was used to test for differences in the timing of VPC between treatments with treatment as a fixed effect and genotype as a random effect. Within each genotype, ANOVA was used to determine if environmental treatment had a significant effect on the time of transition measures, total leaves initiated and leaf initiation rate. For genotypes where treatment had a significant effect (*p* < 0.05), Tukey’s HSD was used to determine which treatments were significantly different from each other. We used one-way ANOVA with natural accession genotypes as the main effect to estimate broad-sense heritability (R^2^ from the ANOVA).

To understand how the timing of VPC impacts plant growth and productivity across environments, we analyzed relationships between the time of transition and reductions in whole plant growth phenotypes of shoot mass, shoot area, and number of leaves initiated in each of the stress environments compared to control. Significant effects of the timing of VPC on these traits was determined using least square linear regression analysis for both accessions and mutant genotypes. Further, we investigated how the timing of VPC impacts plasticity in these traits using a phenotypic plasticity index (PI). This index was calculated as the absolute difference between the maximum and minimum mean values among all four growth conditions divided by the maximum mean value for each genotype (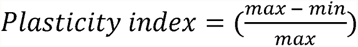), following Valladares *et al*., (2000) and Valladares *et al*., (2006). That is, the PI captured variation in the range of phenotypes observed across environments, normalized by the biggest trait mean among environments. This normalized calculation, giving a value between 0-1, reduces any concern that groups with a greater sample size or mean inappropriately appear to have greater plasticity due to a greater coefficient of variation.

To determine if vegetative phase change contributes to age-dependent changes in phenotypic plasticity, we tested whether developmental phase (i.e. juvenile or adult) impacts plasticity of leaf morphology using PCA and phenotypic plasticity index. Variance of PC1, which described 85% of leaf shape variation, was calculated for juvenile and adult leaves across all four growth environments for mutant and natural accession genotypes to quantify variation in leaf shape. A student’s t-test was used to evaluate whether differences between juvenile and adult plasticity index was significant, where genotypes were the level of observation. Tests were conducted between juvenile and adult leaves of all accessions or across all leaf positions 1-8 for genotypes in the Col-0 background (i.e. miR156/SPL mutants and Col-0 accession) where mutant lines allowed us to examine both juvenile and adult leaves at each position.

To conduct an initial study of potential relationships between plasticity in the timing of vegetative phase change and an accession’s climate of origin, we used the bioclimatic variables from the WorldClim2 data set (Fick & Hijmans, 2017), associated with georeferences for each of the 10 accessions published as part of the 1001 genomes project (Alonso-Blanco *et al*., 2016). We used linear regression to determine any significant relationships between the bioclimatic variables and the phenotypic plasticity index for the time of transition calculated as described above using the number of juvenile leaves.

## Results

### The adult vegetative phase has greater phenotypic plasticity than the juvenile phase

The plasticity of leaf morphology was greater for adult leaves than for juvenile leaves across the four growth environments. Variation in leaf shape between growth environments was also greater in adult leaves (PC1 variance = 0.0075) than in juvenile leaves (PC1 variance = 0.00497) in Col-0 plants with varying levels of miR156/SPL expression, (Fig **1A, B**). Adult leaves also had a higher plasticity index for leaf area, length/width ratio, and circularity, regardless of leaf position (Fig **1C**, Student’s t-test: *p* <0.001). There was no clear difference in leaf shape plasticity between juvenile and adult leaves in natural accessions when PCA was used to describe overall leaf shape (juvenile PC1 variance = 0.005, adult PC1 variance = 0.0041) (Fig **2A, B**). However, in most accessions, adult leaves had greater leaf shape plasticity than juvenile leaves when plasticity was measured by the plasticity index of individual traits,(Fig **2C**). Images of the leaves of these accessions are shown in Figure 3. We conclude that adult leaves are phenotypically more plastic than juvenile leaves.

**Figure 1.**
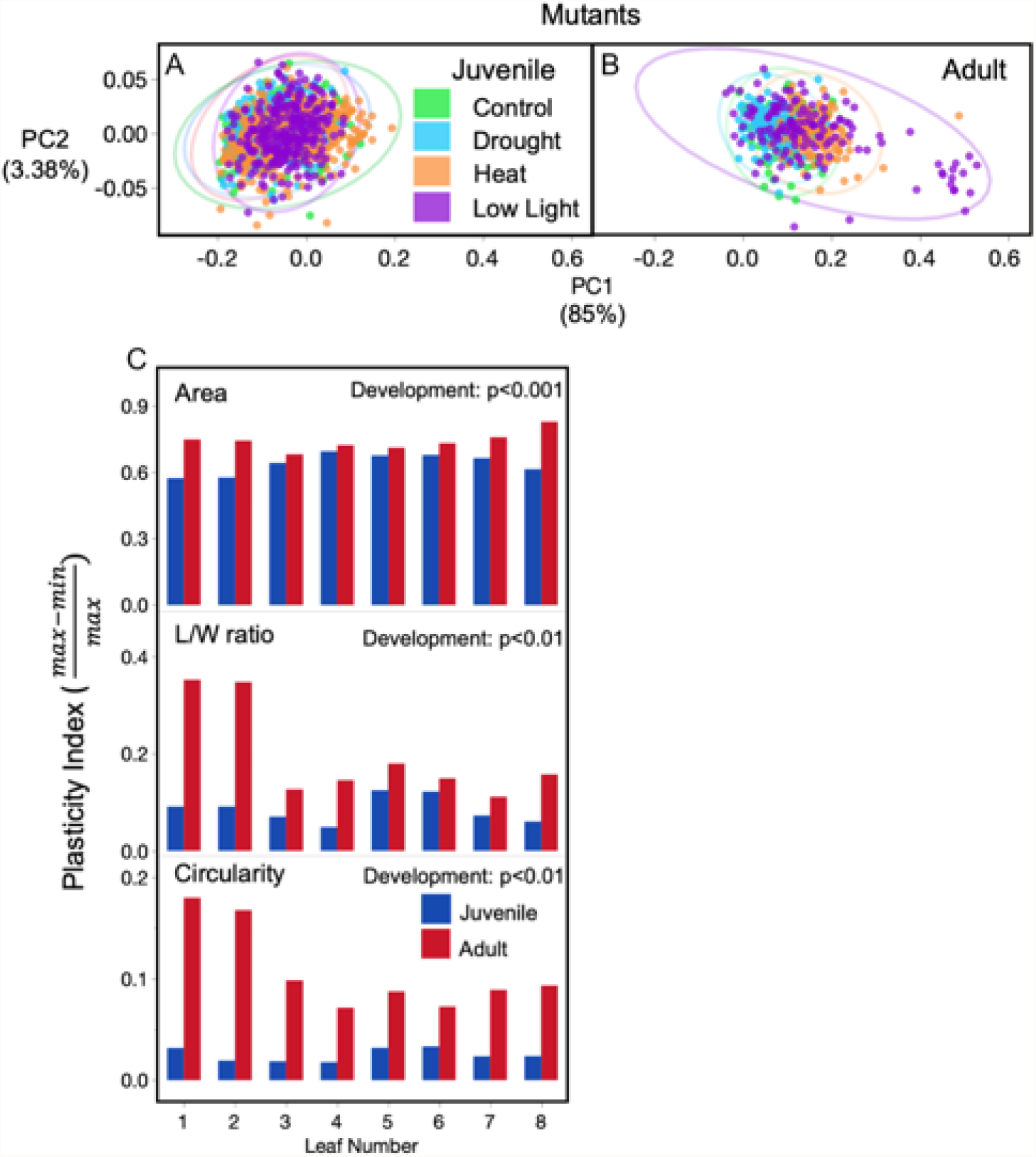
Phenotypic plasticity of juvenile and adult leaf morphology in miR156/SPL mutants. Principal component analysis of leaf shape for juvenile (A) and adult (B) leaves grown in all four environments and phenotypic plasticity index (C) for leaf area, length/width (L/W) ratio, and leaf circularity for adult (red) and juvenile (blue) leaves across leaf positions 1-8. Mutations in the miR156/SPL pathway allow for both juvenile and adult leaves to be produced at each leaf position in the Col-0 genetic background. Significant differences based on Student’s t-test in phenotypic plasticity index between adult and juvenile leaves across all positions denoted in top right corner. A single PCA was conducted on all leaves, but juvenile and adult leaves are plotted separately for ease of visualization.

**Figure 2.**
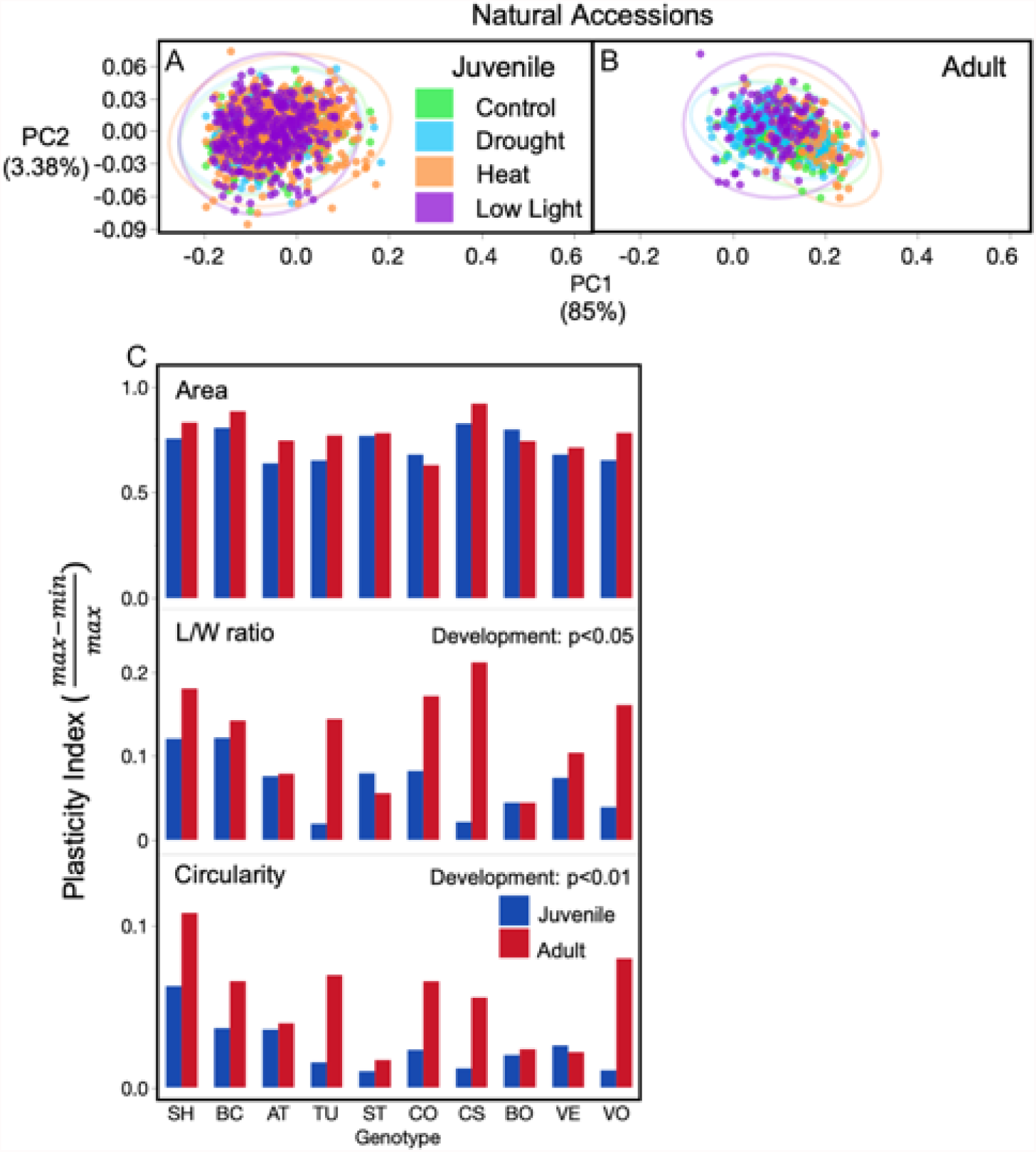
Phenotypic plasticity of juvenile and adult leaf morphology in natural accessions. Principal component analysis of leaf shape for juvenile (A) and adult (B) leaves grown in all four environments and phenotypic plasticity index (C) for leaf area, length/width (L/W) ratio, and leaf circularity for adult (red) and juvenile (blue) leaves of each accession. Significant differences based on Student’s t-test in phenotypic plasticity index between adult and juvenile leaves across all accessions denoted in top right corner. A single PCA was conducted on all leaves, but juvenile and adult leaves are plotted separately for ease of visualization. Full accession names associated with the two-letter codes presented in this figure can be found in Fig.3 and Table S1.

**Figure 3.**
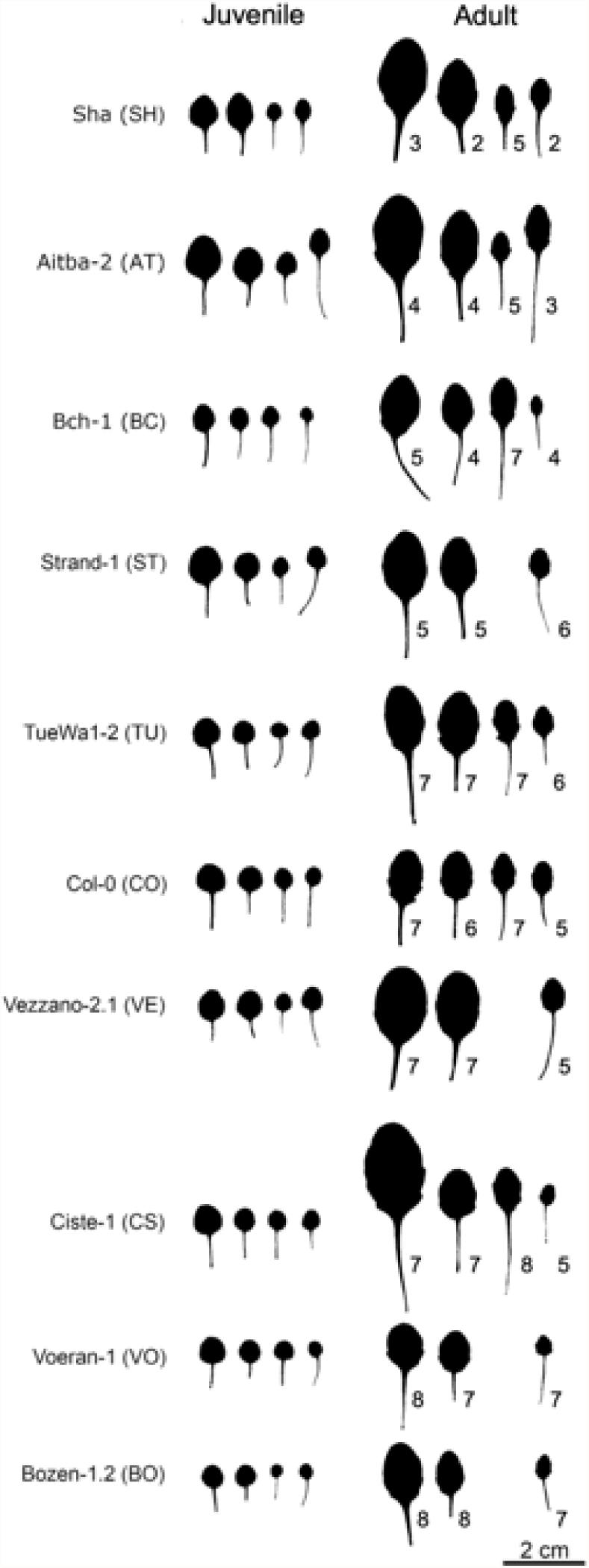
Representative leaf images of juvenile (left) and adult (right) leaves of each accession from control, drought, heat and low light environments shown from left to right. Average position of first adult leaf rounded to the nearest whole number, shown next to adult leaves. Leaves shown are from one of the first two positions of their respective developmental phase.

### The timing of vegetative phase change varies among genotypes and environments

The developmental identity of different rosette leaves was determined using a principal component analysis of phase specific morphological traits. The timing of VPC was measured in two ways: by the number of juvenile leaves, and by the number of days the plant was initiating juvenile leaves, which was calculated using the leaf initiation rate. Both the number of juvenile leaves, and the number of days spent initiating juvenile leaves, differed significantly among genotypes and was altered by abiotic environment (ANOVA: Genotype *p* < 0.0001, Treatment *p* < 0.0001, G x T *p* < 0.0001, GLMM: Days initiating juvenile leaves *p* < 0.0001, number of juvenile leaves *p* < 0.0001) (Fig **4**, Table S3, S4). Differences in the amount of time plants spent in the juvenile phase were largely due to the effect of environmental conditions on the rate of leaf production, as evident from the differences in the total number of leaves produced by 28 days under these conditions (Table S3). Among the accessions examined here, genetic variation led to VPC occurring as early as leaf 3 or day 6, and as late as leaf 8 or day 18 in “control” conditions. VPC in transgenic genotypes with altered miR156 or *SPL* gene expression ranged between leaf 1 or day 1 to leaf 25 or the full 28-day period of the experiment (i.e. these plants never transitioned to adult phase). Broad-sense heritability of the timing of VPC was high, ranging between 0.82 in control and 0.64 in low light environments when measured by the number of juvenile leaves, and 0.61 in drought and 0.46 in control environments, when measured by days initiating juvenile leaves (Table S5).

**Figure 4.**
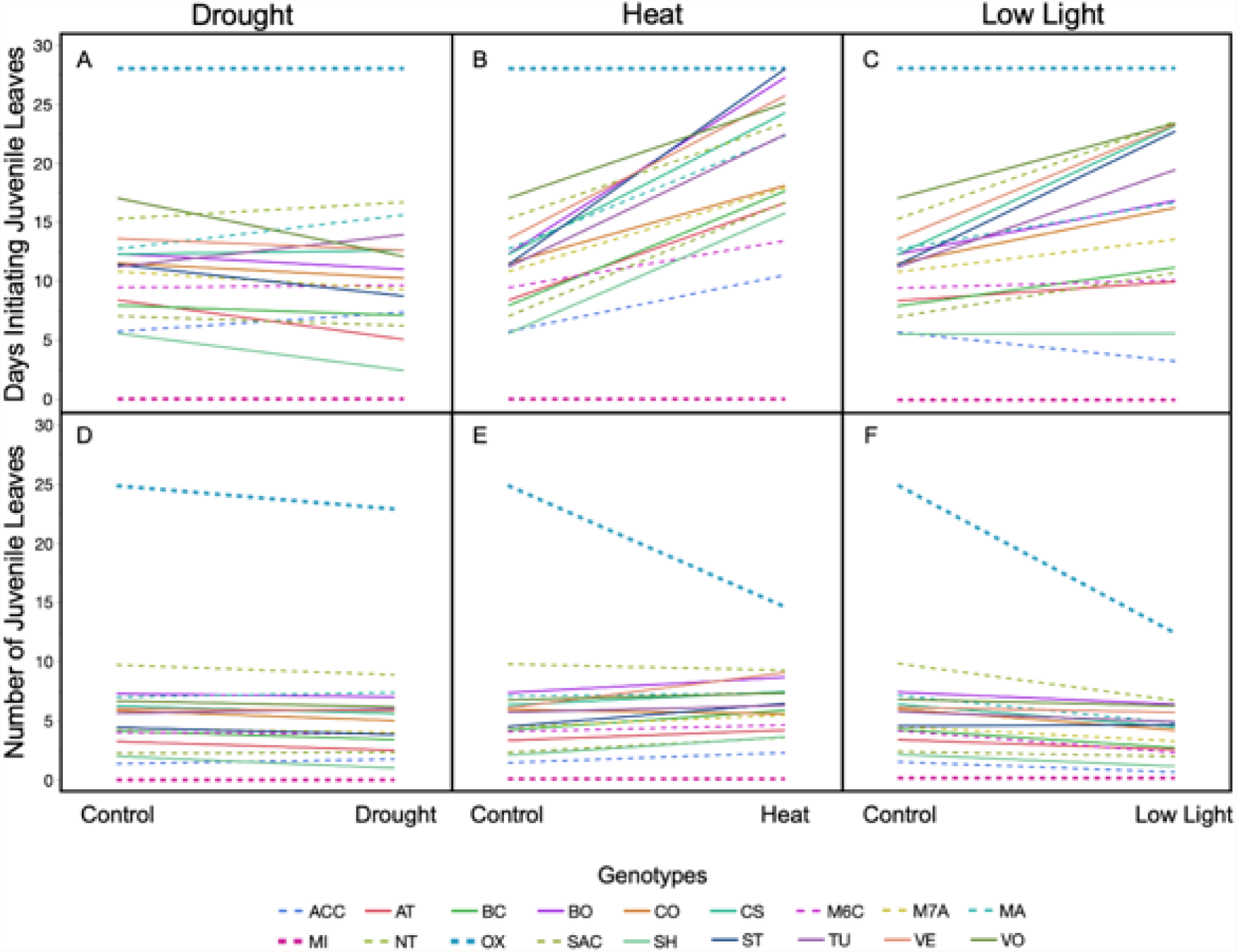
Plasticity in the timing of vegetative phase change, measured as number of days initiating juvenile leaves (A-C) or number of juvenile leaves (D-F), for both natural accession (solid lines) and miR156/SPL mutants (dashed lines) grown in different environments. Each line represents the mean of an individual genotype. The 35S:miR156a overexpression (OX) and MIM:156 (MI) mutants that only produce juvenile or adult leaves respectively are bolded for reference. Full accession names associated with the codes presented in this figure can be found in Table S1.

Plasticity in the timing of VPC in response to abiotic environment differed among genotypes (ANOVA: genotype x treatment = *p* < 0.0001, Fig **4**). Among the natural accessions, heat most often delayed the timing of VPC. Under warm conditions, natural accessions produced, on average, 1.3 additional juvenile leaves compared to control and spent 11.15 additional days initiating juvenile leaves (Table S3). Low light intensity decreased the average number of juvenile leaves, but decreased the rate of leaf initiation by 49%, which was sufficient to increase the number of days most genotypes were in the juvenile phase by an average of 6.39 days. Although most natural accessions produced fewer juvenile leaves and spent less time initiating juvenile leaves in drought, there was no statistically significant differences in the timing of VPC under these conditions compared to control (Table S3). All three environmental treatments subjected plants to some degree of stress as indicated by losses of biomass among all genotypes averaging 8.4, 7.2 and 3 mg in low light, heat and drought respectively.

Differences in the extent and direction of plasticity in the timing of VPC among genotypes in response to abiotic environments led to variation in the order in which genotypes transitioned to the adult phase between environments (Fig. **4**, **S3**). For example, Strand (ST) is the 10^th^ genotype to transition in control, 7^th^ in drought, 17^th^ in heat and 13^th^ in low light when measured by days initiating juvenile leaves. In the MIM156 target mimicry line (MI) and 35S:*MIR156a* (OX) mutants, where miR156 abundance is highly constrained (functionally non-existent or in excess respectively), there was no plasticity in the days spent in the juvenile phase, although the rate of leaf initiation was significantly affected by heat and light intensity in these genotypes.

Our analysis of the plasticity in the timing of VPC in different climatic conditions suggests the response of VPC to environment could contribute to abiotic stress tolerance. Plasticity in the number of juvenile leaves in the natural accessions across all four environments, measured by phenotypic plasticity index, was significantly related to the temperature of the driest quarter (R^2^ = 0.493, *p* < 0.05) and precipitation of the warmest quarter (R^2^=0.502, *p* < 0.05) for each accession’s climate-of-origin (Fig. **S3**). Specifically, increased plasticity in the timing of VPC was related to higher temperatures during the driest quarter and lower precipitation during the warmest quarter.

### The timing of vegetative phase change is correlated with with plant productivity and performance in miR156/SPL mutants, but not in all natural accessions

To understand how changes in the timing of VPC impacts plant productivity and performance in different environments, we explored the relationships between the timing of VPC and the whole plant growth phenotypes of shoot mass, shoot area, and number of leaves initiated during a 28-day period. Among miR156/SPL mutant genotypes, later phase change (i.e. an increase in the number of juvenile leaves or longer time spent initiating juvenile leaves) was significantly associated with a greater reduction in plant growth under environmental stress compared to control (Fig. **5**, **S4**).

**Figure 5.**
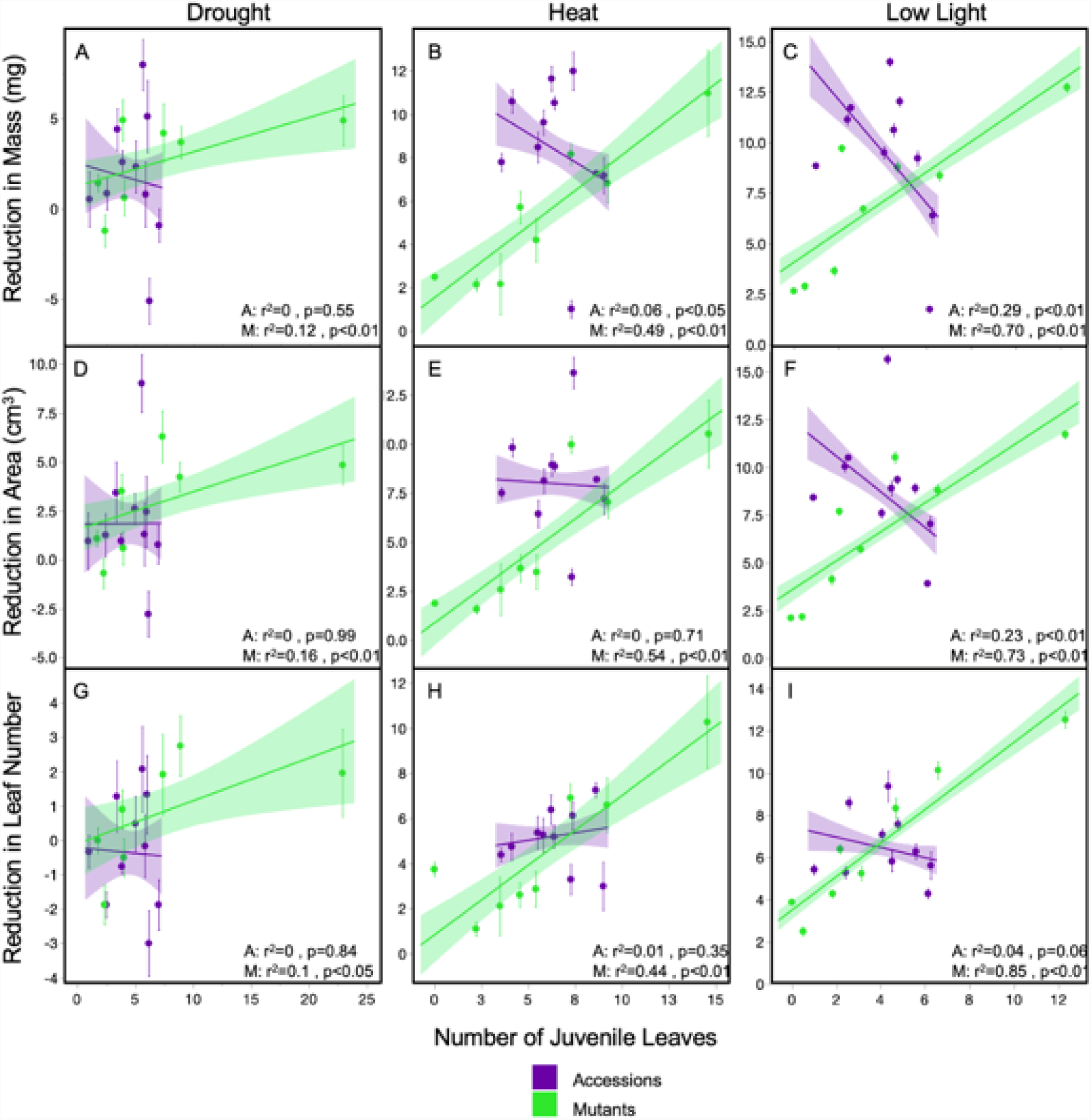
Relationships between the timing of vegetative phase change measured as number of juvenile leaves, and the reduction of whole plant growth phenotypes of shoot mass (A-C), shoot area (D-F), and leaves initiated (G-I) for accessions (purple) and miR156/SPL mutants (green) between control and environmental treatments (drought: A,D,G; heat: B,E,H and low light: C,F,I). *R*^2^ and *p*-values for linear regression of accessions (A) and mutants (M) noted at the bottom of each graph. Points represent the average of each genotype ± standard error.

By contrast, natural genetic variation in the number of juvenile leaves in accessions showed mostly non-significant (*p* > 0.05) or weak (R^2^ < 0.1) relationships with reductions in plant growth in response to each environmental stressor (Fig. **5**). However, plants with a longer juvenile phase, measured as number of juvenile leaves, were better able to maintain shoot biomass and area growth under low light conditions (Fig. **5**, linear regression for number of juvenile leaves vs. reductions in mass: R^2^ = 0.29, *p*<0.01, number of juvenile leaves vs. reductions in area: R^2^ = 0.23, *p*<0.01). Interestingly, some growth responses to low light had contrasting relationships in natural accessions compared to mutant genotypes. Specifically, delayed VPC in the Col-0 background was significantly associated with greater reductions in plant growth in all three stress environments whereas in natural accessions, an earlier transition led to greater reductions in shoot mass and area.

## Discussion

Morphological and physiological plasticity and the timing of development are key ways that organisms cope with fluctuating and unpredictable environments. In plants, there can be plasticity both in the timing of developmental transitions as well as in the character of the organs produced at different times in development (Fig 6). Vegetative phase change (VPC) is a highly conserved developmental transition, but its role in adaptation to the environment is not known. Our study shows that the timing of VPC responds to environmental factors, and demonstrates that this transition alters the amount of phenotypic plasticity that occurs in response to these factors. The existence of natural genetic variation in VPC and its associated plasticity suggests the potential for response to selection, although the VPC-environment-fitness map remains to be dissected.

**Figure 6.**
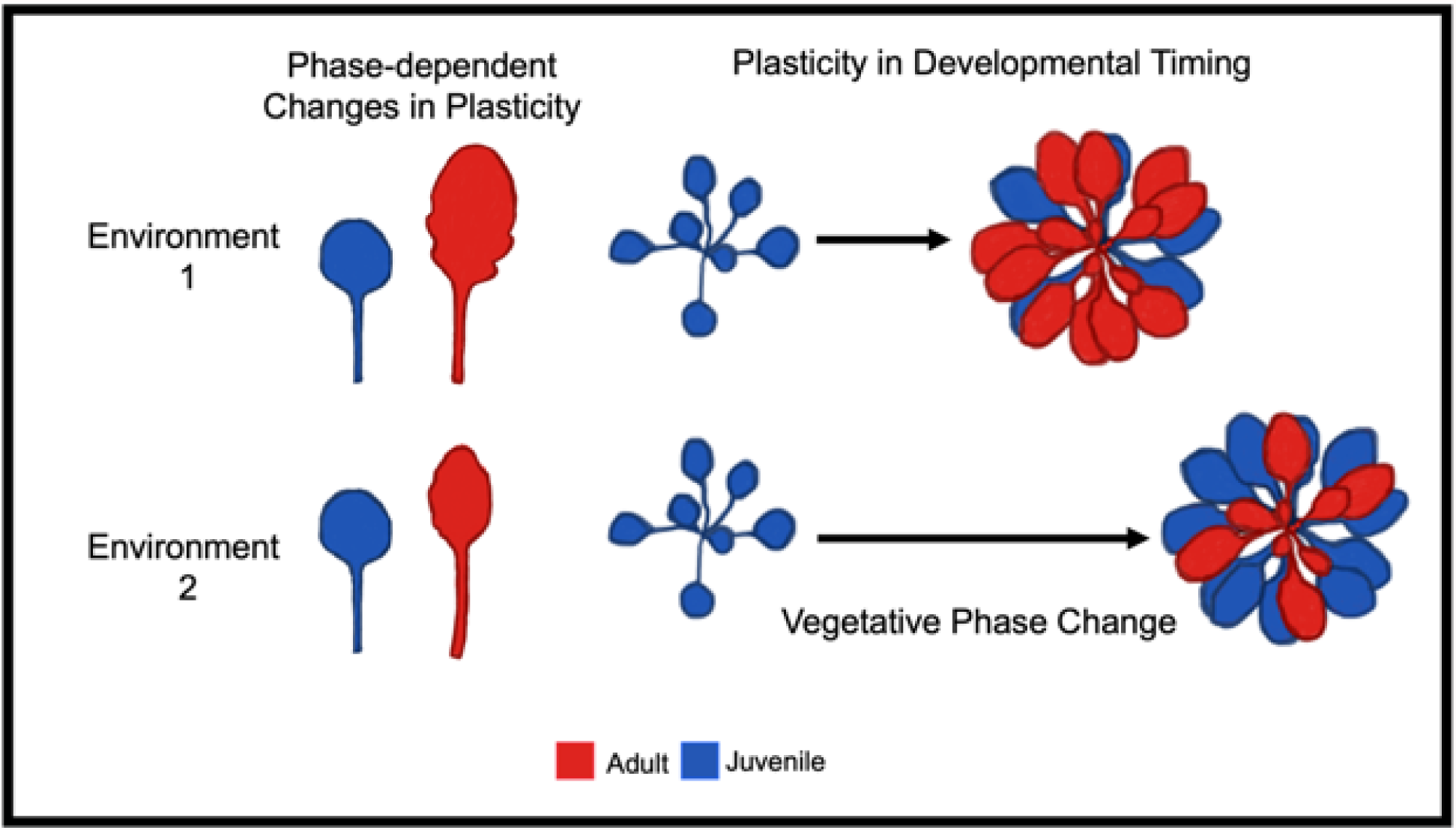
Illustration of two sources of plasticity involving vegetative phase change, phase-dependent changes in plasticity that describe how the developmental phase of an organ (juvenile or adult) impacts the degree of phenotypic plasticity of that organ, and plasticity in developmental timing describing how the timing of vegetative phase change can be accelerated or delayed in response to environment. Similar to what we found, the above example depicts greater plasticity in leaf morphology for adult (red) than juvenile (blue) leaves between environment 1 and 2 as adult leaves change in both shape and size whereas juvenile leaves remain the same. The timing of vegetative phase change is delayed in environment 2 compared to environment 1 as indicated by the longer time arrow and an increased number of juvenile leaves produced.

We found that diverse natural accessions had similar increases in plasticity with vegetative phase change; that is, the juvenile phase was more robust than the adult phase (Fig 6). This consistency suggests this change in plasticity may be adaptive. This increase in plasticity might be favored if it is advantageous to delay high levels of plasticity until after a plant has accumulated a significant amount of information about the environment, but while it still has sufficient time to benefit from a phenotypic adjustment (Fischer *et al*., 2014). Deviations from this pattern, such as in the OX line, which remains in the robust juvenile phase, suggests that there are fitness costs to this strategy, as these OX plants had the greatest reductions in mass, area and leaf initiation in response to environmental stress (Fig **5**). In addition, Fischer *et al*., (2014) predicted the initial more robust phenotype would improve fitness if it adapted a plant to the most likely environment encountered during that life stage. Previous studies show the morphology and physiology of the juvenile phase is well suited to the low light environments often encountered by juvenile leaves emerging in a shaded understory and quickly overtopped by newly initiated leaves (Lawrence *et al*., 2020, 2022; Xu *et al*., 2021), providing support for the hypothesis that VPC modulates phenotypes in an adaptive way.

Changes in phenotypic plasticity modulated by VPC were consistent regardless of genetic background or when the transition occurs. Adult leaves had greater plasticity in leaf shape whether they were produced at node one (i.e. in the MI line) or node ten. This confirms that VPC, specifically expression levels of miR156, is responsible for these changes in plasticity, rather than simply the length of exposure to environmental cues. High levels of miR156, which are associated with a more robust juvenile phase, likely represent an example of microRNA buffering. MicroRNAs buffer by silencing changes in target gene expression (i.e. in response to environmental cues) when target genes are below a certain threshold. Once transcription of the target genes exceeds this threshold, protein output starts to become sensitive to transcriptional changes (Posadas & Carthew, 2014). There are numerous examples of microRNAs contributing to developmental robustness, such as miR164, which buffers its targets and causes robust plant organ development (Sieber *et al*., 2007). The abundance of miR156 early in the juvenile phase far exceeds the level necessary to repress SPL genes, and is therefore likely to contribute to the more robust phenotypes through this buffering mechanism (He *et al*., 2018). Early studies of VPC noted stable phenotypes that were unresponsive to environmental and hormonal treatments in the first two leaves of *Arabidopsis* plants, where miR156 abundance is highest (Telfer *et al*., 1997). An example of miR156 buffering occurs through the microRNA’s role in plant responses to the stress-related phytohormone abscisic acid (ABA). High miR156 levels, as found during the juvenile phase, suppress plant ABA responses through repression of the miR156 target SPL genes, which when expressed during the adult phase, interact directly with the transcription factor ABA-INSENSITIVE5 (ABI5) to facilitate ABA signaling (Dong *et al*., 2021).

Our study found increases in plasticity of leaf shape, a trait that is associated with plant performance, light capture, thermoregulation, and hydraulic efficiency, between the juvenile and adult vegetative phases (Nicotra A *et al*.; Nicotra *et al*., 2011; Adams & Ichiro, 2018; Rowland *et al*., 2020; Strauss *et al*., 2020). Here, adult leaves generally became narrower and more complex with serrations under heat stress and more circular under drought and low light conditions, while juvenile leaves remained similar in shape across environments (Figs **1**, **2**). Although we are unable to say from our data whether these specific shifts in leaf shape of adult leaves confer tolerance to their respective growth environments, previous studies support this hypothesis. Specifically, narrow, and serrated leaves have a reduced boundary layer allowing for more convective cooling but potentially higher rates of water loss and less efficient light absorption, making them better suited for heat, but not low light or drought conditions (Nicotra A *et al*.; Nicotra *et al*., 2011; Adams & Ichiro, 2018; Strauss *et al*., 2020). Many plant traits contribute to environmental tolerance and its possible not all traits will show the same relationships between plasticity and development. For example, in *Iris pumila*, seedling plants have greater plasticity in specific leaf area than older plants in response to changing light environments but chlorophyll content shows the opposite relationship (Avramov *et al*., 2017). It will be important for future studies to include additional phenotypes to understand how changes in plasticity related to development could contribute to adaptation.

Because phenotypic plasticity increases between juvenile and adult phases across genotypes, genetic and environmental variation in the timing of VPC determines when increased plasticity is acquired. We found natural variation in the timing of this transition and significant genotype x environment interactions, indicating the time of VPC has the potential to respond to selection in a manner dependent on environment (i.e. plasticity in developmental timing, Fig 6). We found little plasticity in the timing of VPC in response to drought. But because miR156 expression has previously been shown to respond to drought (Katiyar *et al*., 2015; Liu *et al*., 2019), it is possible that greater plasticity in VPC would occur in a more severe drought treatment. It should also be noted that it is unclear why we observed a decrease in the number of juvenile leaves under low light conditions, given that previous studies have shown that low light intensity increases juvenile leaf number (Leichty & Poethig, 2019; Xu *et al*., 2021). However, we did not carefully control for light quality in these experiments, and it is known that an elevated FR:R ratio can accelerate VPC (Xie *et al*., 2017).

Determining the timing of VPC is not as straightforward as other developmental transitions because it is not marked by any single morphological change, but by multiple traits that do not always appear simultaneously. Because each phase-specific trait can be differentially sensitive to environmental cues, using combinations of traits (as we did) is likely necessary when multiple environments are tested. Despite this increased complexity, understanding how the timing of VPC responds to environments could provide important insights into plant adaptation and acclimation.

Here, we add further evidence that VPC impacts plant growth and productivity as increased levels of miR156 leads to greater vegetative growth across environments, consistent with observations in other species (Fu *et al*., 2012; Rubinelli *et al*., 2013; Wang & Wang, 2015; Zheng *et al*., 2016), while bringing forth new questions about whether these relationships remain true among plants of different genotypes (Fig **5**, Table **S4**). Within the few *Arabidopsis* accessions examined here, we found significant relationships between plasticity in the timing of VPC and multiple climate-of-origin variables (Fig. **S3**), suggesting hypotheses to test in future studies. Further, trait plasticity can influence a plant’s vulnerability to changing climates. For example, plasticity in the leaf traits of 12 perennial plant species was an important determinant of susceptibility to climate change scenarios in a Mongolian steppe (Liancourt *et al*., 2015). Subsequently, climate can alter selection on plasticity. For instance, end-of-season drought conditions selected for increased plasticity of water use efficiency in *A. thaliana* (Kenney *et al*., 2014) whereas plasticity of flowering time in response to temperature was maladaptive (Stinchcombe *et al*., 2004). Modulating age-dependent changes in plasticity might contribute to any role VPC has in adaptation. Additional studies are needed to understand what, if any, type of selection is on VPC and how the timing of vegetative phase change contributes to adaptation and acclimation in plants.

The effects of VPC on plasticity found here in *A. thaliana* may be conserved across species. VPC is regulated by miR156 and its sister microRNAs across all land plants, allowing for microRNA buffering to create a consistently more robust juvenile phase. For example, across 40 species of *Passiflora,* juvenile leaf morphology is conserved despite the highly variable morphology among adult leaves (Chitwood & Otoni, 2017), similar to what we observed here between the various genotypes (Fig. **1**, **2**). This further suggests the age-dependent changes in plasticity due to VPC could lead to varying degrees of both intra and interspecific trait variation through time among populations. It seems likely many published studies capture patterns of variation driven by VPC within and among species, although authors may have overlooked this transition (i.e. (Mason *et al*., 2013; Garbowski *et al*., 2021). Intraspecific variation can play an important role in community- and ecosystem-level processes (Bolnick *et al*., 2011; Madritch *et al*., 2014; Turner *et al*., 2020; Westerband *et al*., 2021), though the functional basis of this variation is usually not understood. To better understand the implications of VPC on these large scale processes, further characterization of the timing of VPC across plant species (currently only done in a few; maize, (Poethig, 1988; Bongard-Pierce *et al*., 1996); *Arabidopsis thaliana*, (Telfer *et al*., 1997); *Eucalyptus globulus,* (Jordan *et al*., 1999); *Nicotiana tabacum*, (Feng *et al*., 2016); Sorghum, (Hashimoto *et al*., 2019); Acacia, (Leichty & Poethig, 2019); *Passiflora edulis*, (Silva *et al*., 2019); Poplar, (Lawrence *et al*., 2021) and Rice, (Asai *et al*., 2002)) and variation among genotypes within species (even fewer; *Eucalyptus globulus*, (Jordan *et al*., 1999); Maize, (Poethig, 1988; Foerster *et al*., 2015); and *A. thaliana*, (Doody *et al*., 2022)) is necessary.

VPC may contribute to plant adaptation by setting up periods of low and high phenotypic plasticity when it is most beneficial. Here we demonstrate that the timing of vegetative phase change varies across genotypes, interacts with the environment, and alters the plasticity of vegetative traits. While more work is needed to fully understand the functional, ecological, and evolutionary significance of VPC, its determination as a modulator of age-dependent changes in plasticity provides new insights for understanding plant environmental interactions.

## Acknowledgments

We thank Erin Doody for providing preliminary data regarding the timing of vegetative phase change in *Arabidopsis thaliana* accessions. This research was funded by NSF grants DGE-1845298 and NSF IOS-2109780 awarded to EHLP, NSF DEB-1927009 and NIH R35GM138300 grants awarded to JRL, and NIH GM51893 awarded to RSP.

## Author Contributions

EHLP, RSP and JRL contributed to research planning and design, EHLP performed the experiments and statistical analysis, EHLP wrote the initial version of the manuscript and all authors revised and provided comments.

## Supplemental Information

**Table S1** *A. thaliana* accessions and mutants and their and associated IDs

**Table S2** Environmental factors for each growth treatment

**Table S3** Time of VPC and leaf initiation across environments

**Table S4** Statistical tests on timing of VPC

**Table S5** Broad-sense heritability for the timing of vegetative phase change

**Figure S1** Leaf shape outlines and PCA morphospace

**Figure S2** Determination of the timing of VPC using PCA

**Figure S3** Plasticity index for timing of VPC for each genotype

**Figure S4** Plasticity in VPC related to climate of origin

**Figure S5** Relationships between the timing of VPC measured as days initiating juvenile leaves and reductions in whole plant growth phenotypes from environmental stress

## Notes

### Competing Interest Statement

The authors have declared no competing interest.

### Summary of Updates

We have reanalyzed data, used more rigorous statists and made significant improvements to the text

## References

Adams WW, Ichiro III. 2018. The Leaf: A Platform for Performing Photosynthesis.

Alonso-Blanco C, Andrade J, Becker C, Bemm F, Bergelson J, Borgwardt KM, Cao J, Chae E, Dezwaan TM, Ding W, et al. 2016. 1,135 Genomes reveal the global pattern of polymorphism in *Arabidopsis thaliana*. Cell 166: 481–491.

Arshad M, Feyissa BA, Amyot L, Aung B, Hannoufa A. 2017. MicroRNA156 improves drought stress tolerance in alfalfa (*Medicago sativa*) by silencing SPL13. Plant Science 258: 122–136.

Arshad M, Gruber MY, Hannoufa A. 2018. Transcriptome analysis of microRNA156 overexpression alfalfa roots under drought stress. Scientific Reports 8: 9363.

Asai K, Satoh N, Sasaki H, Satoh H, Nagato Y. 2002. A rice heterochronic mutant, mori1, is defective in the juvenile-adult phase change. Development 129: 265–273.

Avramov S, Miljković D, Klisarić NB, Živković U, Tarasjev A. 2017. Ontogenetic plasticity of anatomical and ecophysiological traits and their correlations in Iris pumila plants grown in contrasting light conditions. Plant Species Biology 32: 392–402.

Barton KE. 2008. Phenotypic plasticity in seedling defense strategies: compensatory growth and chemical induction. Oikos 117: 917–925.

Bassiri A, Irish EE, Scott PR. 1992. Heterochronic effects of Teopod 2 on the growth and photosensitivity of the maize shoot. The Plant Cell 4: 497–504.

Blázquez MA, Ahn JH, Weigel D. 2003. A thermosensory pathway controlling flowering time in Arabidopsis thaliana. Nature genetics 33: 168–171.

Bolnick DI, Amarasekare P, Araújo MS, Bürger R, Levine JM, Novak M, Rudolf VHW, Schreiber SJ, Urban MC, Vasseur DA. 2011. Why intraspecific trait variation matters in community ecology. Trends in Ecology & Evolution 26: 183–192.

Bongard-Pierce DK, Evans MMS, Poethig RS. 1996. Heteroblastic features of leaf anatomy in maize and their genetic regulation. International Journal of Plant Sciences 157: 331.

Bonhomme V, Picq S, Gaucherel C, Claude J. 2014. Momocs: Outline analysis using R. Journal of Statistical Software 56: 1–24.

Chien JC, Sussex IM. 1996. Differential regulation of trichome formation on the adaxial and abaxial leaf surfaces by gibberellins and photoperiod in *Arabidopsis thaliana* (L.) Heynh. Plant Physiology 111: 1321–1328.

Chitwood DH, Otoni WC. 2017. Morphometric analysis of *Passiflora* leaves: the relationship between landmarks of the vasculature and elliptical Fourier descriptors of the blade. GigaScience 6: 1–13.

Dong H, Yan S, Jing Y, Yang R, Zhang Y, Zhou Y, Zhu Y, Sun J. 2021. MIR156-Targeted SPL9 is phosphorylated by SnRK2s and interacts with ABI5 to enhance ABA responses in *Arabidopsis*. Frontiers in Plant Science 12: 708573.

Doody E, Zha Y, He J, Poethig RS. 2022. The genetic basis of natural variation in the timing of vegetative phase change in Arabidopsis thaliana. Development 149.

Dorken ME, Barrett SCH. 2004. Phenotypic plasticity of vegetative and reproductive traits in monoecious and dioecious populations of *Sagittaria latifolia* (*Alismataceae*): a clonal aquatic plant. Journal of Ecology 92: 32–44.

Feng S, Xu Y, Guo C, Zheng J, Zhou B, Zhang Y, Ding Y, Zhang L, Zhu Z, Wang H, et al. 2016. Modulation of miR156 to identify traits associated with vegetative phase change in tobacco (*Nicotiana tabacum*). Journal of Experimental Botany 67: 1493–1504.

Fick SE, Hijmans RJ. 2017. WorldClim 2: new 1-km spatial resolution climate surfaces for global land areas. International Journal of Climatology 37: 4302–4315.

Fischer B, van Doorn GS, Dieckmann U, Taborsky B. 2014. The evolution of age-dependent plasticity. American Naturalist 183: 108–125.

Foerster JM, Beissinger T, de Leon N, Kaeppler S. 2015. Large effect QTL explain natural phenotypic variation for the developmental timing of vegetative phase change in maize (*Zea mays L*.). Theoretical and Applied Genetics 128: 529–538.

Forster MA, Bonser SP. 2009. Heteroblastic development and shade-avoidance in response to blue and red light signals in acacia implexa. Photochemistry and Photobiology 85: 1375–1383.

Franco-Zorrilla JM, Valli A, Todesco M, Mateos I, Puga MI, Rubio-Somoza I, Leyva A, Weigel D, García JA, Paz-Ares J, et al. 2007. Target mimicry provides a new mechanism for regulation of microRNA activity. Nature Genetics 39: 1033–1037.

Fu C, Sunkar R, Zhou C, Shen H, Zhang J-YY, Matts J, Wolf J, Mann DGJ, Stewart CN, Tang Y, et al. 2012. Overexpression of miR156 in switchgrass (*Panicum virgatum* L.) results in various morphological alterations and leads to improved biomass production. Plant Biotechnology Journal 10: 443–452.

Garbowski M, Johnston DB, Brown CS. 2021. Leaf and root traits, but not relationships among traits, vary with ontogeny in seedlings. Plant and Soil.

Ghalambor CK, McKay JK, Carroll SP, Reznick DN. 2007. Adaptive versus non-adaptive phenotypic plasticity and the potential for contemporary adaptation in new environments. Functional Ecology 21: 394–407.

Hashimoto S, Tezuka T, Yokoi S. 2019. Morphological changes during juvenile-to-adult phase transition in *sorghum*. Planta 250.

He J, Xu M, Willmann MR, McCormick K, Hu T, Yang L, Starker CG, Voytas DF, Meyers BC, Poethig RS. 2018. Threshold-dependent repression of SPL gene expression by miR156/miR157 controls vegetative phase change in *Arabidopsis thaliana*. PLOS Genetics 14: e1007337.

Hoverman JT, Relyea RA. 2007. How felxible is phenotypic plasticity? Developmental windows for trait induction and reversal. Ecology 88: 693–705.

Jordan GJ, Potts BM, Wiltshire RJE. 1999. Strong, independent, quantitative genetic control of the timing of vegetative phase change and first flowering in *Eucalyptus globulus* ssp. *globulus* (Tasmanian Blue Gum). Heredity 83: 179–187.

Katiyar A, Smita S, Muthusamy SK, Chinnusamy V, Pandey DM, Bansal KC. 2015. Identification of novel drought-responsive microRNAs and trans-acting siRNAs from *Sorghum bicolor (L.)* Moench by high-throughput sequencing analysis. Frontiers in Plant Science 6: 506.

Kenney AM, McKay JK, Richards JH, Juenger TE. 2014. Direct and indirect selection on flowering time, water-use efficiency (WUE, δ13C), and WUE plasticity to drought in *Arabidopsis thaliana*. Ecology and Evolution 4: 4505–4521.

Kleunen M Van, Fischer M. 2005. Constraints on the evolution of adaptive phenotypic plasticity in plants. New Phytologist 166: 49–60.

Kong Y, Elling A, Chen B, Deng XW. 2010. Differential expression of microRNAs in maize inbred and hybrid lines during salt and drought stress. American Journal of Plant Sciences 01: 69–76.

Lawrence EH, Leichty AR, Doody EE, Ma C, Strauss SH, Poethig RS. 2021. Vegetative phase change in *Populus tremula x alba*. New Phytologist 231: nph.17316.

Lawrence EH, Springer CJ, Helliker BR, Poethig RS. 2020. MicroRNA156-mediated changes in leaf composition lead to altered photosynthetic traits during vegetative phase change. New Phytologist: nph.17007.

Lawrence EH, Springer CJ, Helliker BR, Poethig RS. 2022. The carbon economics of vegetative phase change. Plant Cell and Environment 45: 1286–1297.

Lee H, Yoo SJ, Lee JH, Kim W, Yoo SK, Fitzgerald H, Carrington JC, Ahn JH. 2010. Genetic framework for flowering-time regulation by ambient temperature-responsive miRNAs in *Arabidopsis*. Nucleic acids research 38: 3081–93.

Leichty AR, Poethig RS. 2019. Development and evolution of age-dependent defenses in ant-acacias. Proceedings of the National Academy of Sciences 116: 15596–15601.

Levy YY, Dean C. 1998. Control of flowering time. Current Opinion in Plant Biology 1: 49–54.

Liancourt P, Boldgiv B, Song DS, Spence LA, Helliker BR, Petraitis PS, Casper BB. 2015. Leaf-trait plasticity and species vulnerability to climate change in a Mongolian steppe. Global Change Biology 21: 3489–3498.

Liu X, Zhang X, Sun B, Hao L, Liu C, Zhang D, Tang H, Li C, Li Y, Shi Y, et al. 2019. Genome-wide identification and comparative analysis of drought-related microRNAs in two maize inbred lines with contrasting drought tolerance by deep sequencing. PLoS ONE 7: e0219176.

Madritch MD, Kingdon CC, Singh A, Mock KE, Lindroth RL, Townsend PA. 2014. Imaging spectroscopy links aspen genotype with below-ground processes at landscape scales. Philosophical Transactions of the Royal Society B: Biological Sciences 369.

Mason CM, Mcgaughey SE, Donovan LA. 2013. Ontogeny strongly and differentially alters leaf economic and other key traits in three diverse *Helianthus* species. Journal of Experimental Botany 64: 4089–4099.

May P, Liao W, Wu Y, Shuai B, Richard McCombie W, Zhang MQ, Liu QA. 2013. The effects of carbon dioxide and temperature on microRNA expression in *Arabidopsis* development. Nature Communications 4: 2145.

Meuthen D, Baldauf SA, Bakker TCM, Thünken T. 2018. Neglected patterns of variation in phenotypic plasticity: age-and sex-specific antipredator plasticity in a cichlid fish. The America Naturalist 191: 475–490.

Munir J, Dorn LA, Donohue K, Schmitt J. 2001. The effect of maternal photoperiod on seasonal dormancy in *Arabidopsis thaliana* (*Brassicaceae*). American Journal of Botany 88: 1240–1249.

Nicotra A AB, Leigh AB, Kevin Boyce CC, Jones D CS, Niklas E KJ, Royer F DL, Tsukaya HG. The evolution and functional signicance of leaf shape in the angiosperms.

Nicotra AB, Leigh A, Boyce CK, Jones CS, Niklas KJ, Royer DL, Tsukaya H. 2011. The evolution and functional significance of leaf shape in the angiosperms. Functional Plant Biology 38: 535–552.

Niinemets Ü. 2016. Leaf age dependent changes in within-canopy variation in leaf functional traits: a meta-analysis. Journal of Plant Research 129: 313–338.

Nilsson-Örtman V, Rogell B, Stoks R, Johansson F. 2015. Ontogenetic changes in genetic variances of age-dependent plasticity along a latitudinal gradient. Heredity 2015 115:4 115: 366–378.

Olszewski N, Sun T, Gubler F. 2002. Gibberellin signaling biosynthesis, catabolism, and response pathways. The Plant Cell 14: S61–S80.

Parrish JAD, Bazzaz FA. 1985. Ontogenetic niche shifts in old-field annuals. Ecology 66: 1296–1302.

Poethig RS. 1988. Heterochronic mutations affecting shoot development in maize. Genetics 119: 959–73.

Poethig RS. 1990. Phase change and the regulation of shoot morphogenesis in plants. Science 250: 923–930.

Posadas DM, Carthew RW. 2014. MicroRNAs and their roles in developmental canalization. Current Opinion in Genetics and Development 27: 1–6.

Rose KME, Mickelbart M V, Jacobs DF. 2019. Plasticity of phenotype and heteroblasty in contrasting populations of Acacia koa. Annals of Botany 124: 399–409.

Rowland SD, Zumstein K, Nakayama H, Cheng Z, Flores AM, Chitwood DH, Maloof JN, Sinha NR. 2020. Leaf shape is a predictor of fruit quality and cultivar performance in tomato. New Phytologist 226: 851–865.

Rubinelli PM, Chuck G, Li X, Meilan R. 2013. Constitutive expression of the *Corngrass1* microRNA in poplar affects plant architecture and stem lignin content and composition. Biomass and Bioenergy 54: 312–321.

Sebestyén F, Miklós M, Iván K, Tökölyi J. 2020. Age-dependent plasticity in reproductive investment, regeneration capacity and survival in a partially clonal animal (*Hydra oligactis*). Journal of Animal Ecology 89: 2246–2257.

Sieber P, Wellmer F, Gheyselinck J, Riechmann JL, Meyerowitz EM. 2007. Redundancy and specialization among plant microRNAs: role of the MIR164 family in developmental robustness. Development 134: 1051–1060.

Silva PO, Batista DS, Henrique J, Cavalcanti F, Koehler AD, Vieira LM, Fernandes AM, Hernan Barrera-Rojas C, Ribeiro DM, Nogueira FTS, et al. 2019. Leaf heteroblasty in *Passiflora edulis* as revealed by metabolic profiling and expression analyses of the microRNAs miR156 and miR172. Annals of Botany 123: 1191–1203.

Stinchcombe JR, Dorn LA, Schmitt J. 2004. Flowering time plasticity in Arabidopsis thaliana: a reanalysis of Westerman & Lawrence (1970). Journal of Evolutionary Biology 17: 197–207.

Strauss S, Lempe J, Prusinkiewicz P, Tsiantis M, Smith RS. 2020. Phyllotaxis: is the golden angle optimal for light capture? New Phytologist 225: 499–510.

Telfer A, Bollman KM, Poethig RS. 1997. Phase change and the regulation of trichome distribution in *Arabidopsis thaliana*. Development 124: 645–654.

Tsukaya H, Shoda K, Kim G-T, Uchimiya H. 2000. Heteroblasty in *Arabidopsis thaliana* (L.) Heynh. Planta 2000 210:4 210: 536–542.

Turner KG, Lorts CM, Haile AT, Lasky JR. 2020. Effects of genomic and functional diversity on stand-level productivity and performance of non-native *Arabidopsis*. Proceedings of the Royal Society B 287: 20202041.

Valladares F, Sanchez-Gomez D, Zavala MA. 2006. Quantitative estimation of phenotypic plasticity: bridging the gap between the evolutionary concept and its ecological applications. Journal of Ecology 94: 1103–1116.

Valladares F, Wright SJ, Lasso E, Kitajima K, Pearcy RW. 2000. Plastic phenotypic response to light of 16 congeneric shrubs from a panamanian rainforest. Ecology 81: 1925–1936.

Wang JW, Park MY, Wang LJ, Koo Y, Chen XY, Weigel D, Poethig RS. 2011. MiRNA control of vegetative phase change in trees. PLoS Genetics 7: e1002012.

Wang H, Wang H. 2015. The miR156/SPL module, a regulatory hub and versatile toolbox, gears up crops for enhanced agronomic traits. Molecular Plant 8: 677–688.

Westerband AC, Funk JL, Barton KE. 2021. Intraspecific trait variation in plants: a renewed focus on its role in ecological processes. Annals of botany 127: 397–410.

Willmann MR, Poethig RS. 2007. Conservation and evolution of miRNA regulatory programs in plant development. Current Opinion in Plant Biology 10: 503–511.

Wu G, Park MY, Conway SR, Wang JW, Weigel D, Poethig RS. 2009. The sequential action of miR156 and miR172 regulates developmental timing in *Arabidopsis*. Cell 138: 750–759.

Wu G, Poethig RS. 2006. Temporal regulation of shoot development in *Arabidopsis thaliana* by miR156 and its target SPL3. Development 133: 3539–3547.

Xie Y, Liu Y, Wang H, Ma X, Wang B, Wu G, Wang H. 2017. Phytochrome-interacting factors directly suppress MIR156 expression to enhance shade-avoidance syndrome in Arabidopsis. Nature Communications 2017 8:1 8: 1–11.

Xu M, Hu T, Poethig RS. 2021. Low light intensity delays vegetative phase change. Plant Physiology kiab243.

Xu M, Hu T, Zhao J, Park M-Y, Earley KW, Wu G, Yang L, Poethig RS. 2016. Developmental functions of miR156-regulated SQUAMOSA PROMOTER BINDING PROTEIN-LIKE (SPL) genes in *Arabidopsis thaliana*. PLOS Genetics 12: e1006263.

Yan A, Pan J, An L, Gan Y, Feng H. 2012. The responses of trichome mutants to enhanced ultraviolet-B radiation in *Arabidopsis thaliana*. Journal of Photochemistry and Photobiology B: Biology 113: 29–35.

Yang L, Conway SR, Poethig RS. 2011. Vegetative phase change is mediated by a leaf-derived signal that represses the transcription of miR156. Development 138: 245–249.

Yang L, Xu M, Koo Y, He J, Poethig RS. 2013. Sugar promotes vegetative phase change in *Arabidopsis thaliana* by repressing the expression of MIR156A and MIR156C. eLife 2: e00260.

Zhang J, Lechowicz MJ. 1994. Correlation between time of flowering and phenotypic plasticity in *Arabidopsis thaliana* (*Brassicaceae*). American Journal of Botany 81: 1336–1342.

Zhao J, Doody E, Poethig RS. 2023. Reproductive competence is regulated independently of vegetative phase change in Arabidopsis thaliana. Current Biology 33: 487–497.

Zheng Q-L, Liu J, Goff BM, Dinkins RD, Zhu H. 2016. Genetic manipulation of miR156 for improvement of biomass production and forage quality in Red Clover. Crop Science 56: 1–7.

