## Supplemental Information for "Vegetative Phase Change Causes Age-Dependent Changes in Phenotypic Plasticity"

*New Phytologist Supporting Information*

Article acceptance date: N/A

The following supporting information is available for this article:

**Table S1**. *A. thaliana* accessions and the miR156/SPL mutants in the Col-0 background used in this study and their and associated IDs

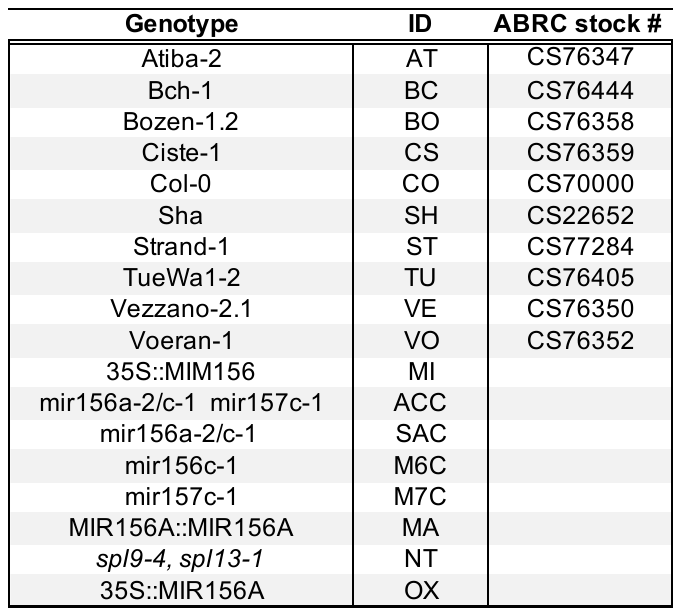

**Table S2**. Environmental factors for each growth treatment.

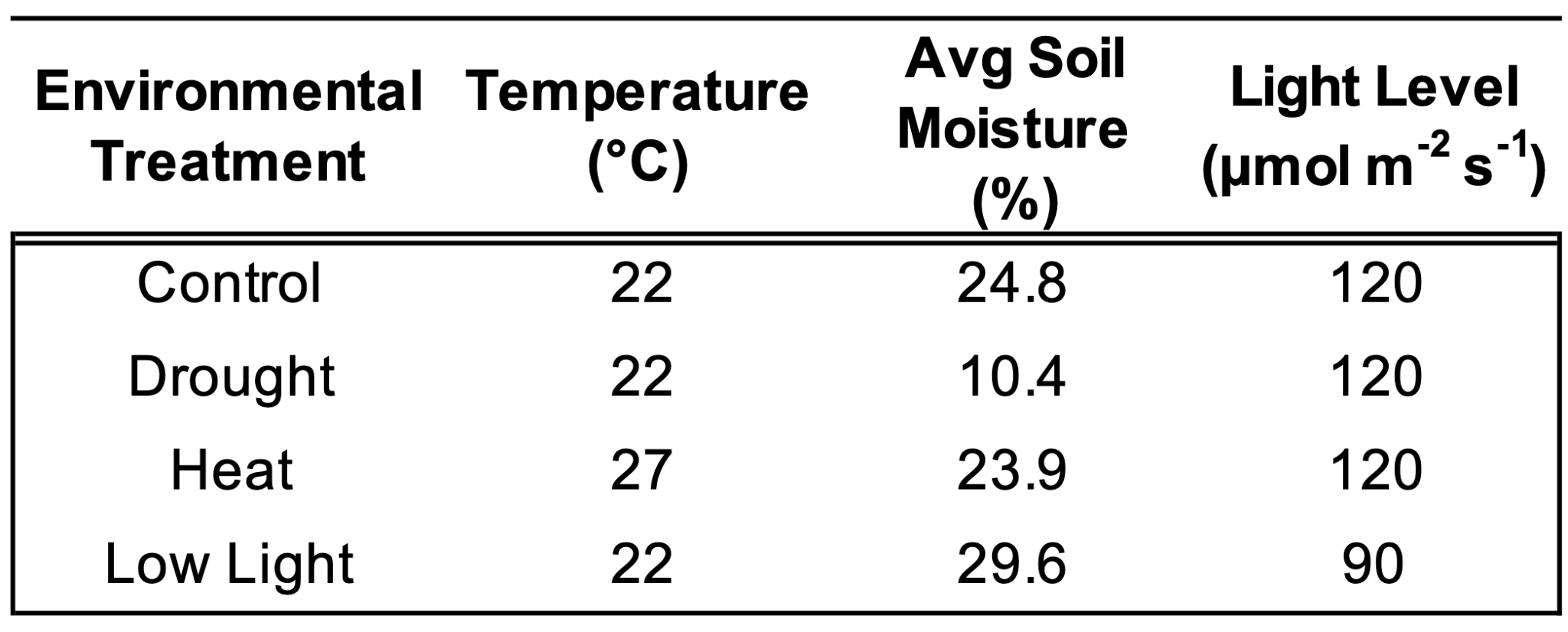

**Table S3**. Timing of vegetative phase change, measured as number of juvenile leaves produced or the number of days initiating juvenile leaves, and leaf initiation traits during 28-days of growth for natural accessions and miR156/SPL mutants grown in different environments. Values represent the mean ± standard error. Different lowercase letters indicate treatment groups that are significantly different (*p* < 0.05) based on Tukey’s HSD, following a significant ANOVA result (*p* < 0.05). Groups significantly different from control are bolded. N represents the number of plants sampled.

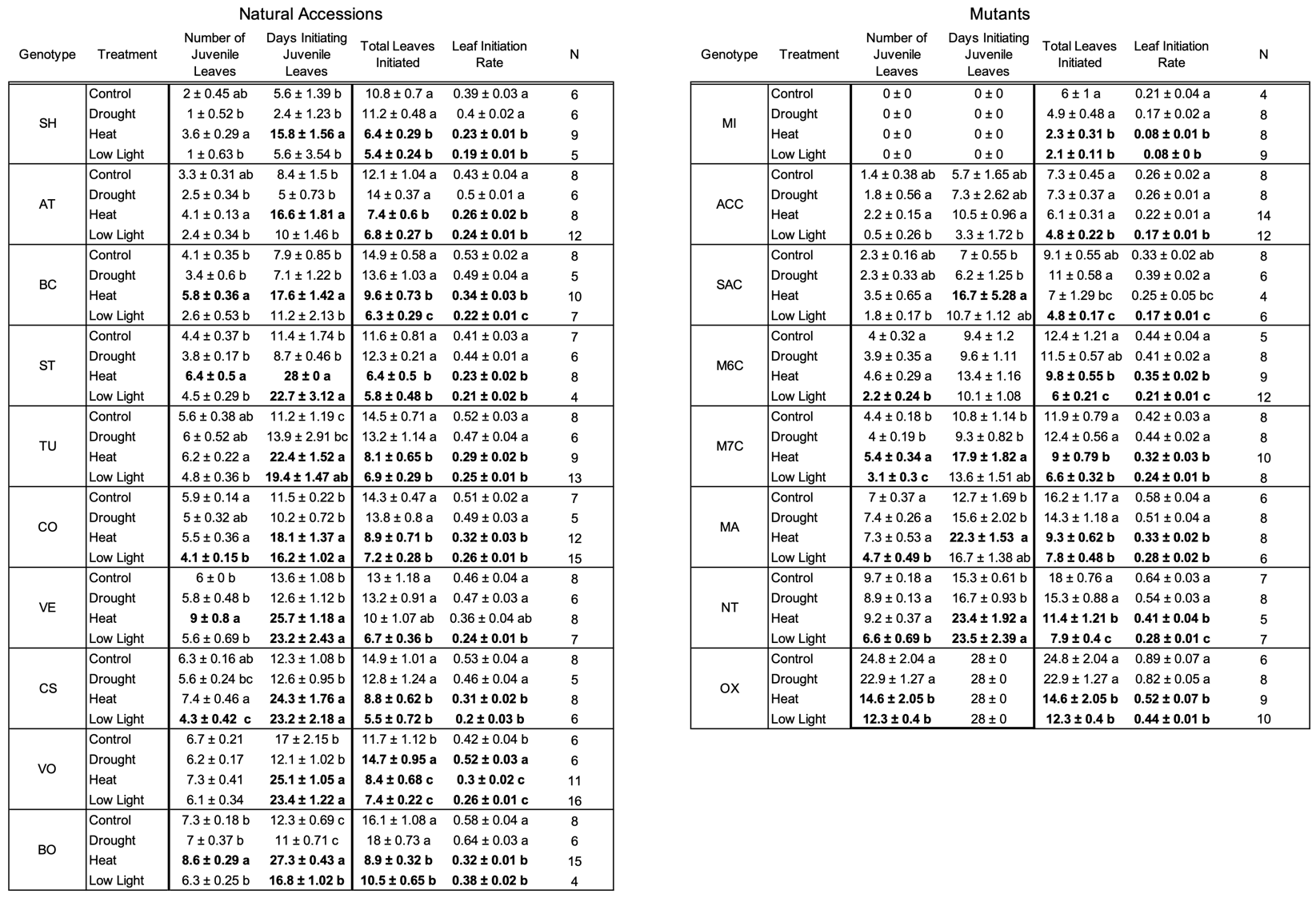

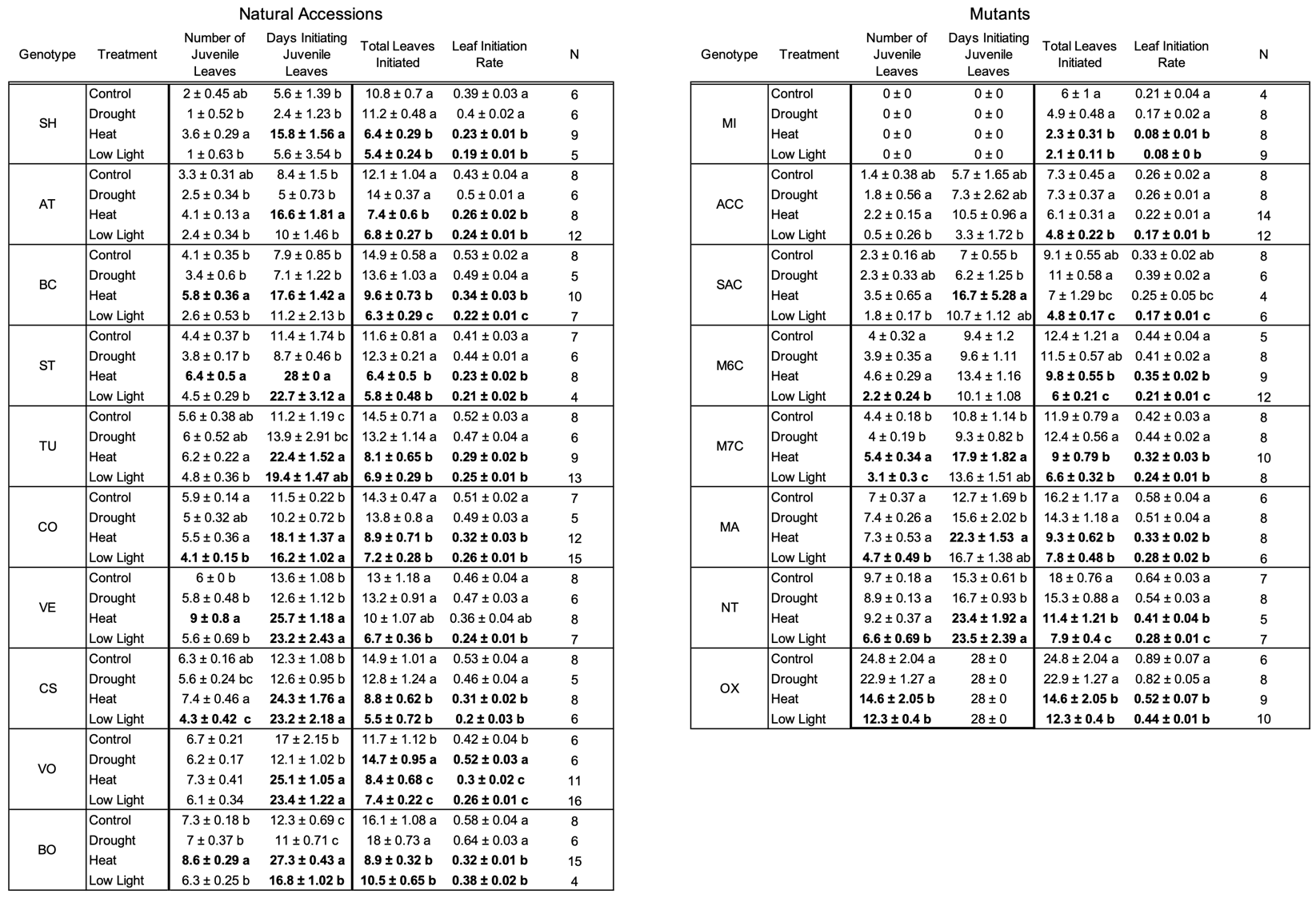

**Table S4.** Statistical tests on timing of vegetative phase change measured as number of juvenile leaves and days initiating juvenile leaves for plants grown in different treatment environments (control heat, drought, and low light)

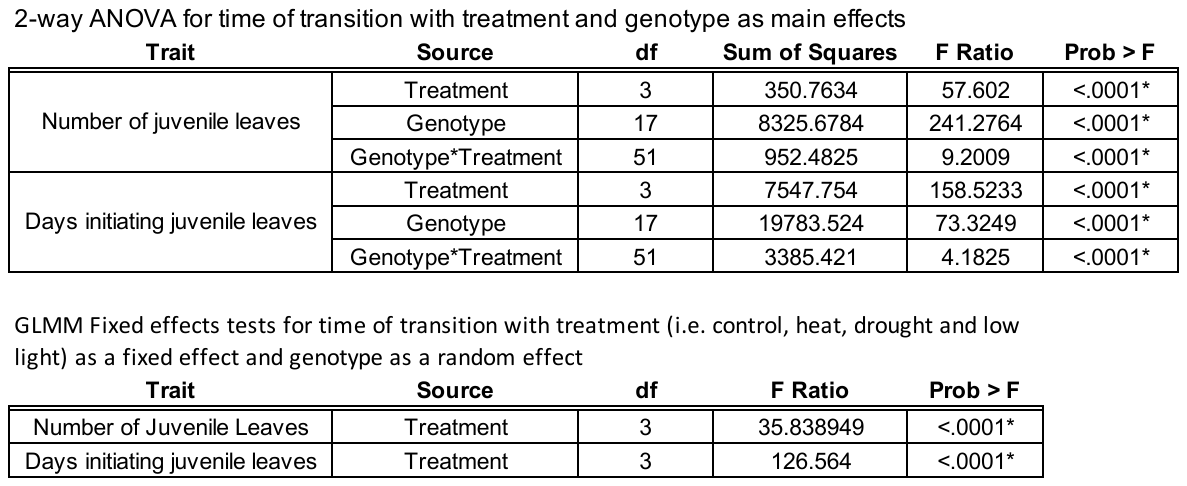

**Table S5.** Broad-sense heritability for the timing of vegetative phase change in *Arabidopsis thaliana* in each treatment environment measured by the number of juvenile leaves or days initiating juvenile leaves. Estimates determined by one-way ANOVA with natural accession genotypes as the main effect. RMSE represents root mean square error.

| **Broad Sense Heritability for the Timing of VPC** | | | | |
| --- | --- | --- | --- | --- |
| **Treatment** | **Juvenile Leaves** | | **Days** | |
|  | **R^2^** | **RMSE** | **R^2^** | **RMSE** |
| Control | 0.82 | 0.77 | 0.46 | 3.39 |
| Drought | 0.82 | 0.9 | 0.61 | 2.65 |
| Heat | 0.66 | 1.25 | 0.58 | 3.95 |
| Low Light | 0.64 | 1.2 | 0.57 | 5.22 |

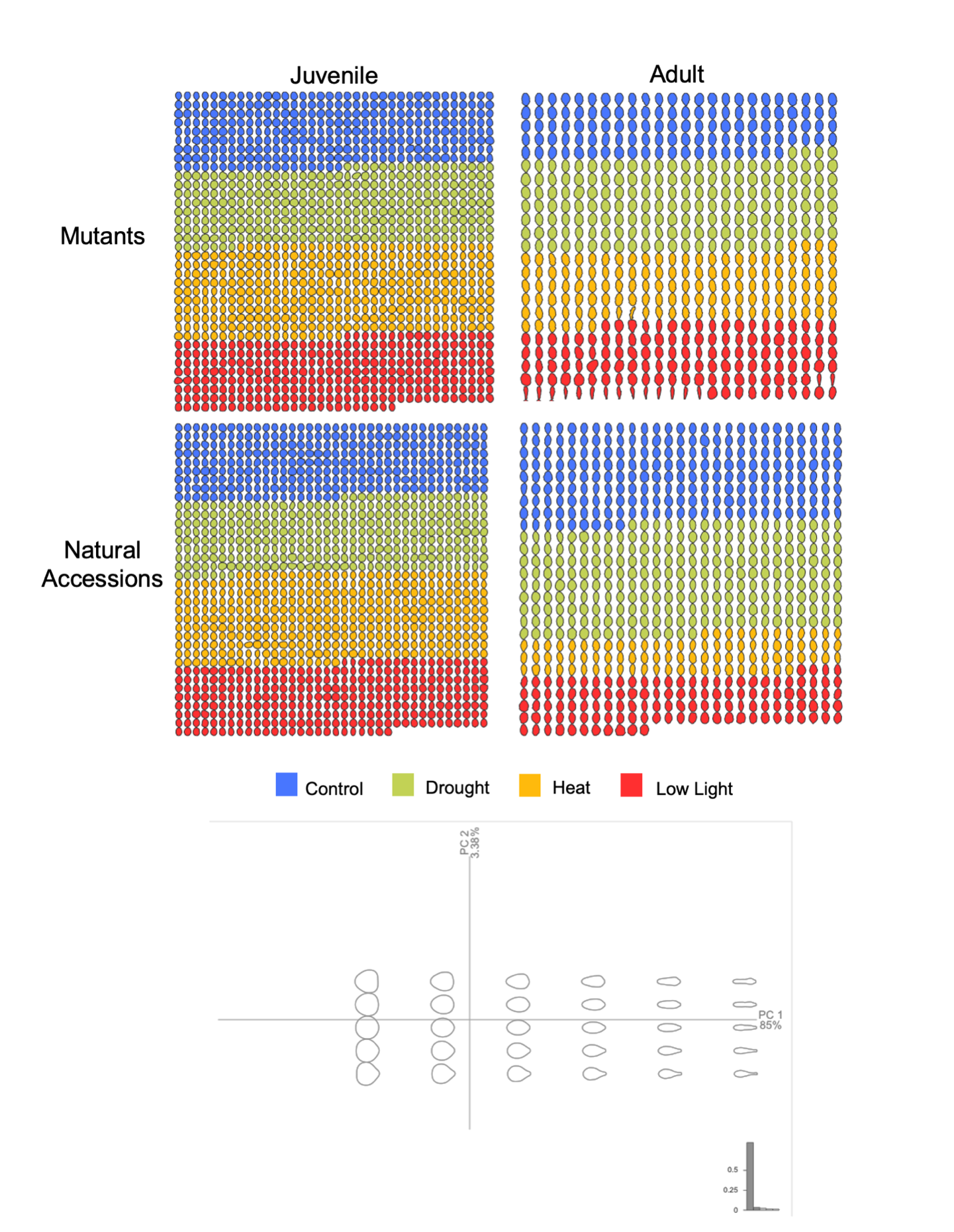

**Figure S1.** Outlines of leaf shapes used in Momocs R package morphometrics analysis color coded by growth environment and the PCA morphospace showing how leaf shape changes along PC1 and PC2 axes.

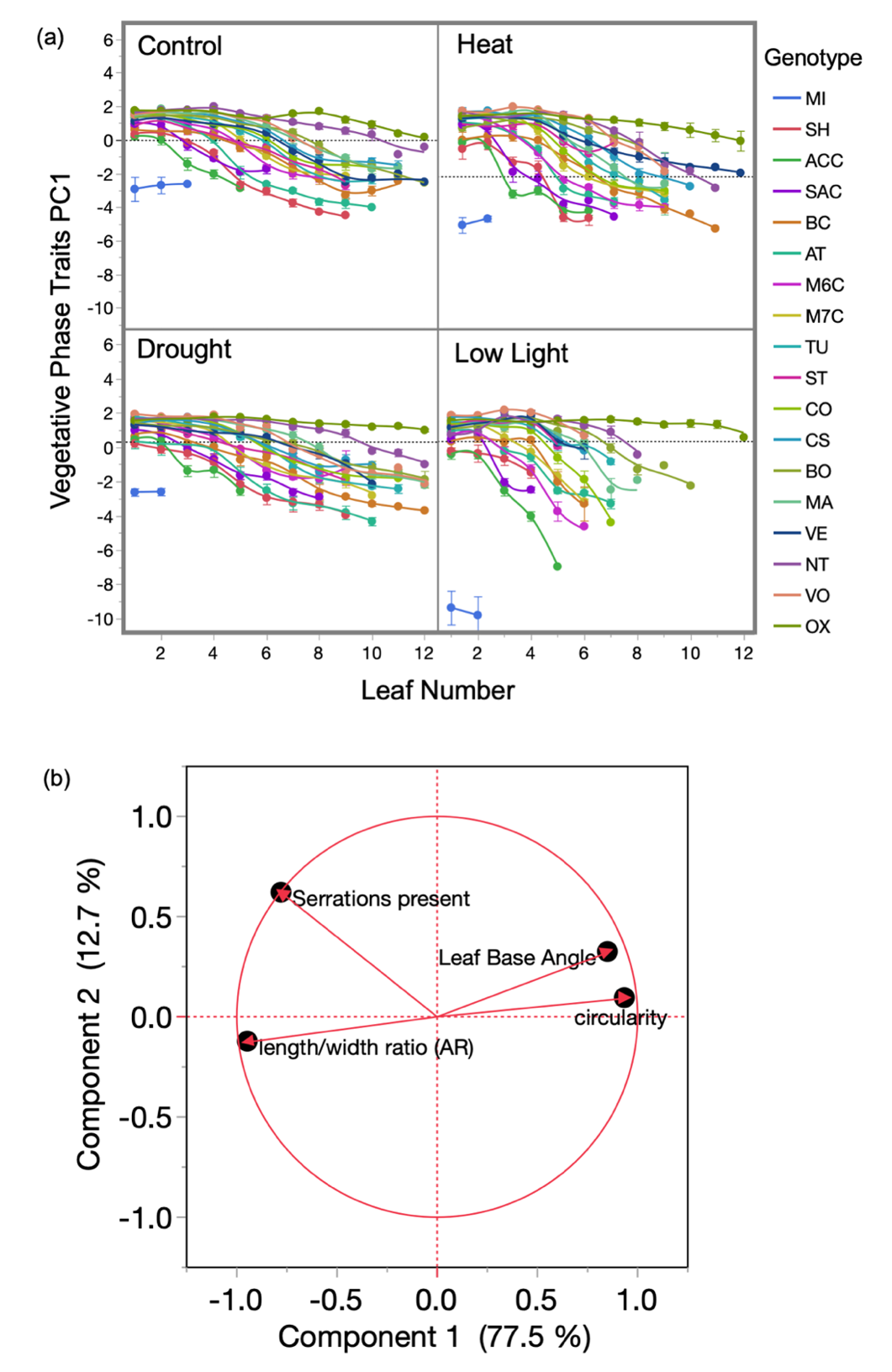

**Figure S2**. Determination of the timing of vegetative phase change using a PC1 summarizing vegetative phase specific traits of serrations, leaf length/width ratio, leaf base angle, and circularity. Vegetative phase traits PC1 for each genotype across leaf positions (1 = first leaf produced) in control, heat, drought and low light treatments (a). Dashed grey horizontal line indicates the PC1 value below all leaves of 35S:miR156a overexpression mutants (OX), which only produce juvenile leaves, used to determine the onset of adult morphology. Loading plot of vegetative phase specific traits (b).

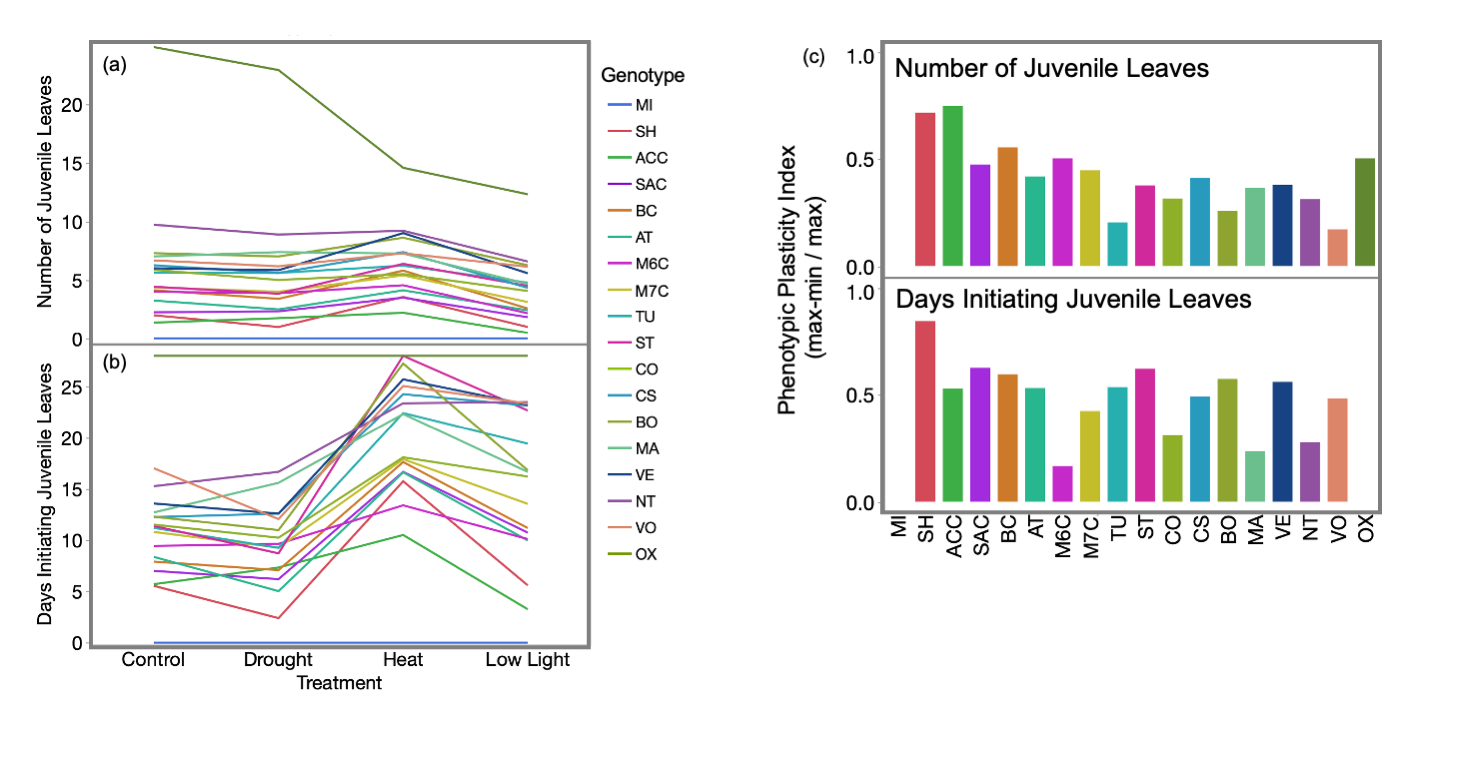

**Figure S3.** Phenotypic plasticity index over all four environments for time of transition measures for each genotype. Genotypes are ordered from earliest to latest transition times based on number of juvenile leaves produced in control conditions.

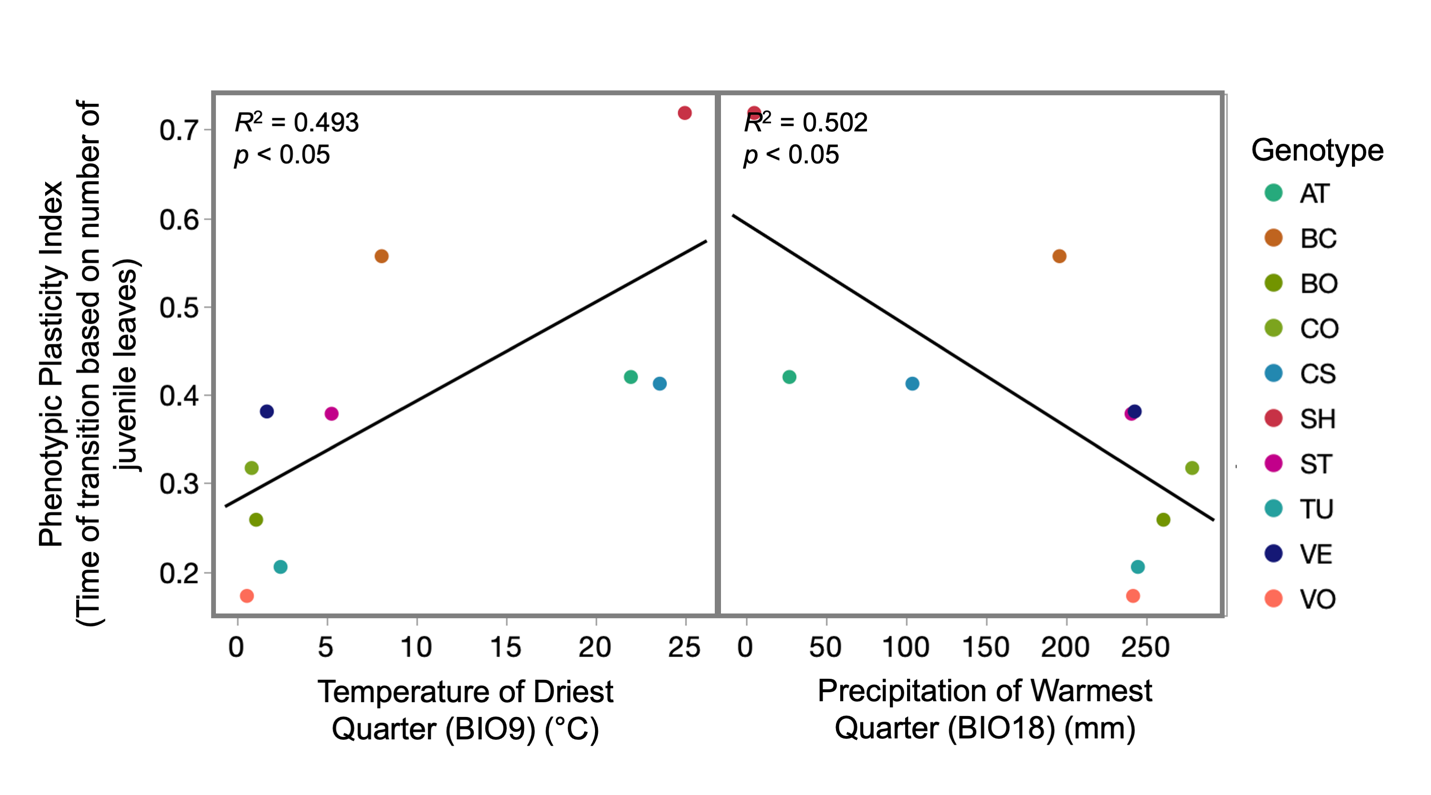

**Figure S4.** Plasticity in the timing of vegetative phase change is related to climate of origin across studied accessions. Phenotypic plasticity index across the four tested environments for timing of vegetative phase change measured as number of juvenile leaves produced is positively associated with temperature of the driest quarter (WorldClim BIO9) (left) and negatively related to precipitation of warmest quarter (WorldClim BIO18) (right) of the accession’s climate of origin. *R*^2^ and *p*-values for the linear regression denoted in the top left of each graph.

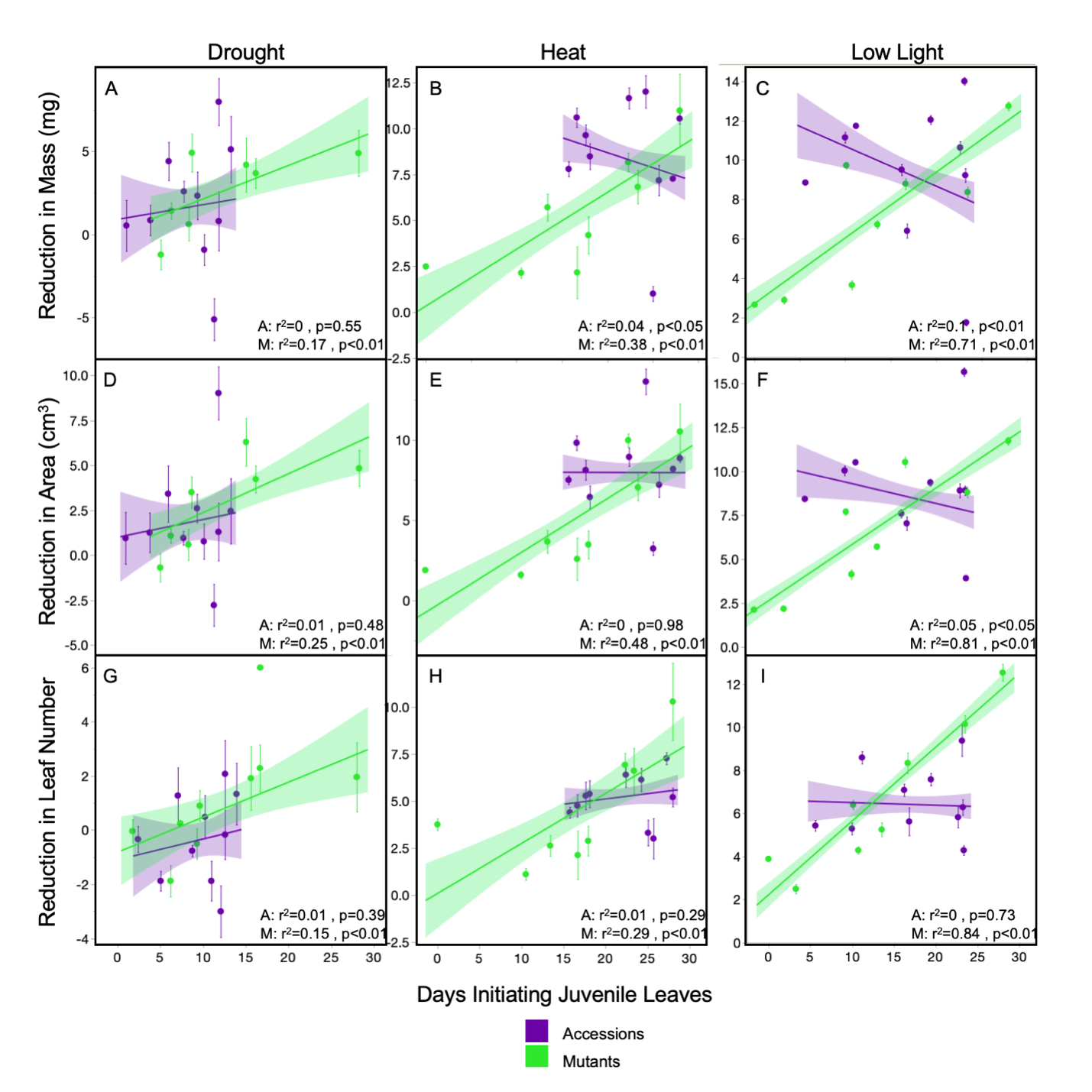

**Figure S5**. Relationships between the timing of vegetative phase change measured as days initiating juvenile leaves, and the reduction of whole plant growth phenotypes of shoot mass (A-C), shoot area (D-F), and leaves initiated (G-I) for accessions (purple) and miR156/SPL mutants (green) between control and environmental treatments (drought: A,D,G; heat: B,E,H and low light: C,F,I). *R*^2^ and *p*-values for linear regression of accessions (A) and mutants (M) noted at the bottom of each graph. Points represent the average of each genotype ± standard error.
